# The Manganese Salt (MnJ) Functions as A Potent Universal Adjuvant

**DOI:** 10.1101/783910

**Authors:** Rui Zhang, Chenguang Wang, Yukun Guan, Xiaoming Wei, Mengyin Sha, Miao Jing, Mengze Lv, Jing Xu, Yi Wan, Zhengfan Jiang

**Author notes:** These authors contributed equally (Rui Zhang, Chenguang Wang, Yukun Guan).

## Abstract

Aluminum adjuvants have been used for a century in various vaccines due to its ability to potentiate humoral immunity and safety records since 1920s. Manganese is an essential micronutrient required for diverse biological activities in cells. We previously found that Mn^2+^ is a strong type I-interferon stimulator activating the cGAS-STING pathway. Herein we report that a colloidal manganese salt (MnJ) is a potent adjuvant to induce both humoral and cellular immune responses, particularly CTL activation. When administrated intranasally, MnJ was also a strong mucosal adjuvant, inducing high levels of IgA antibodies. MnJ strongly promoted dendritic cell maturation and antigen-specific T cell activation. Interestingly, IL-1/-18 induction and release by Mn^2+^-activated ASC-mediated inflammasomes were not observed. MnJ showed great adjuvant effects to all tested antigens including inactivated viruses, recombinant proteins and peptides by either intramuscular or intranasal immunization. These findings may have implications in developing potent but safe Mn^2+^-containing vaccines.

## Main Text

Vaccination is one of the most successful public health interventions. Adjuvants are widely used to increase the immunogenicity of vaccines with various advantages: (1) reducing the amount of antigens; (2) reducing the number of immunizations; (3) inducing a faster protection; (4) improving the efficacy of vaccines in newborns, the elderly or immunocompromised populations ^1, 2^. Over the past few decades, large amounts of adjuvants have been developed. According to their different mechanisms of action, adjuvants are divided into two categories: immune potentiators and delivery systems. Some immune potentiators, composed of pathogen-associated molecular patterns (PAMPs) or synthesized activators of pattern-recognition receptors (PPRs), activate innate immune responses to induce the subsequent production of cytokines and chemokines. Other potentiators, composed of cells or cytokines like dendritic cells, IL-12 or GM-CSF, directly activate immunity. Delivery systems including liposome, micelles, virosome, nanoparticles, microsphere, Oil/Water emulsion, virus-like particles (VLP) and immune stimulating complexes (ISCOM), usually carry antigen to target cells and assist antigen uptake by antigen presenting cells (APCs) ^3, 4^.

The immune enhancement effect of aluminum salts (Alum) was first reported by Glenny et al. in the 1920’s when they found that injection of guinea pigs with diphtheria toxoid precipitated with potassium aluminum provided greater protection than toxoid alone ^5^. Since then, Alum-containing adjuvants have been employed in billions of doses of vaccines and administered annually to millions of people ^6^. In fact, Alum is the only human adjuvant widely used, partly due to its minimal reactogenicity and inexpensiveness ^7^. However, Alum-adjuvanted vaccines need repeated administrations which may cause or manifest its adverse effects including increased IgE production and neurotoxicity ^8^. In addition, Alum mainly induces T helper 2 (TH2) cell response through NLPR3 inflammasome activation ^9–11^, but not TH1 or cytotoxic T-lymphocyte (CTL) response ^1, 12^. Therefore, Alum is generally believed to be unable to elicit cellular immune responses that are essential for virus or tumor vaccines.

In the past decades, however, the medical need for new adjuvants is increasing as (1) the tremendously increased use of purified antigens like recombinant proteins with low immunogenicity due to the absence of immunostimulatory components recognized by PRRs; (2) adjuvants inducing cellular immune responses especially CTL are badly needed for virus and cancer vaccines ^13–15^. So far few adjuvants have been approved by the US Food and Drug Administration for use in humans and several formulations are in clinical trials. The oil-in-water MF59 in influenza vaccines for the elderly was approved in the 1990’s, followed by AS03 in vaccines against avian influenza virus, AS04 in hepatitis B virus (HBV) and human papillomavirus (HPV) vaccines, and AS01 in herpes zoster virus vaccines ^16^. Toll-like receptor (TLR) agonists, like the CpG DNA and poly (I:C), have been studied in the past two decades as new adjuvant candidates ^7^. cGAS-STING ^17–20^ agonists like DMXAA ^21^, c-di-GMP ^22^, cGAMP ^23^ and Chitosan ^24^ also showed some adjuvant effects.

Manganese (Mn) is a nutritional inorganic trace element required for a variety of physiological processes including development, reproduction, neuronal function and antioxidant defenses ^25, 26^. Mn (Mn^2+^ in general cases) is essential for some metalloenzymes such as Mn superoxide dismutase (SOD2, Mn^3+^ or Mn^2+^ in this case), glutamine synthetase (GS), and arginase ^27^. However, its function in regulating immunity is largely unknown. Previously we found that Mn^2+^ was required for the host defense against DNA virus by increasing the sensitivity of the DNA sensor cGAS and its downstream adaptor protein STING. Mn^2+^ was released from mitochondria and Golgi apparatus upon virus infection and accumulated in the cytosol where it bound directly to cGAS, enhancing the sensitivity of cGAS to double-stranded DNA (dsDNA) and its enzymatic activity. Mn^2+^ also enhanced cGAMP-STING binding affinity ^28^. Importantly, Mn^2+^ was a potent innate immune stimulator, inducing type I-IFN and cytokine production in the absence of any infection.

Herein we report that Manganese (Mn^2+^) functions as a universal non-inflammatory adjuvant to induce both humoral and cellular immune responses, particularly CTL activation. A colloidal manganese salt (MnJ) was found to strongly promote dendritic cell maturation and antigen-specific T cell activation. When administrated intranasally, MnJ was also a strong mucosal adjuvant, inducing high levels of IgA antibodies. MnJ showed great adjuvant effects to all tested antigens including inactivated viruses, recombinant proteins and peptides by either intramuscular or intranasal immunization. Thus, we identified the second metal element that functions as adjuvant nearly one hundred years after Alum was found.

## Results

### Mn^2+^ Promotes DC Maturation via cGAS-STING Activation

We previously found that Mn^2+^ is a strong type I-IFN stimulator by activating the cGAS-STING pathway in the absence of any infections *in vitro* and *in vivo*, we thus hypothesized that Mn^2+^ would have some adjuvant potentials. Since APCs play an important role in linking innate and adaptive immunity, we first determined whether Mn^2+^ promotes DC maturation. RNA-seq analysis on Mn^2+^- or LPS-treated mouse bone marrow derived dendritic cells (BMDCs) revealed that Mn^2+^ induced robust production of both IFNβ and various IFNαs, which were not induced by LPS (Fig. 1a), together with significantly up-regulated costimulatory molecules CD80 and CD86, mouse MHC-I proteins H-2K/D/Q, immunoproteasome subunits PSMB8/9, peptide transporters TAP1/2 and chemokines, including CCL2 and CCL3, that increase the recruitment of immune cells to the injection site. Surprisingly, compared to LPS-treated BMDCs, Mn^2+^-treated BMDCs did not produce pro-inflammatory cytokines including IL-1α/β and IL-18, nor IL-10 or IL-12, suggesting that Mn^2+^ triggered a distinct signaling in BMDCs, which were confirmed by qPCR analysis (Fig. 1b). Additionally, Mn^2+^-induced expression of costimulatory molecules CD86, CD80 and CD40 were lost in BMDCs from *Tmem173*^−⁄−^, *Irf3^−⁄−^Irf7*^−⁄−^ and *Ifnar*^−⁄−^ mice, showing that Mn^2+^-induced DC maturation was completely dependent on the activation of the cGAS-STING pathway (Fig. 1c).

**Fig. 1.**
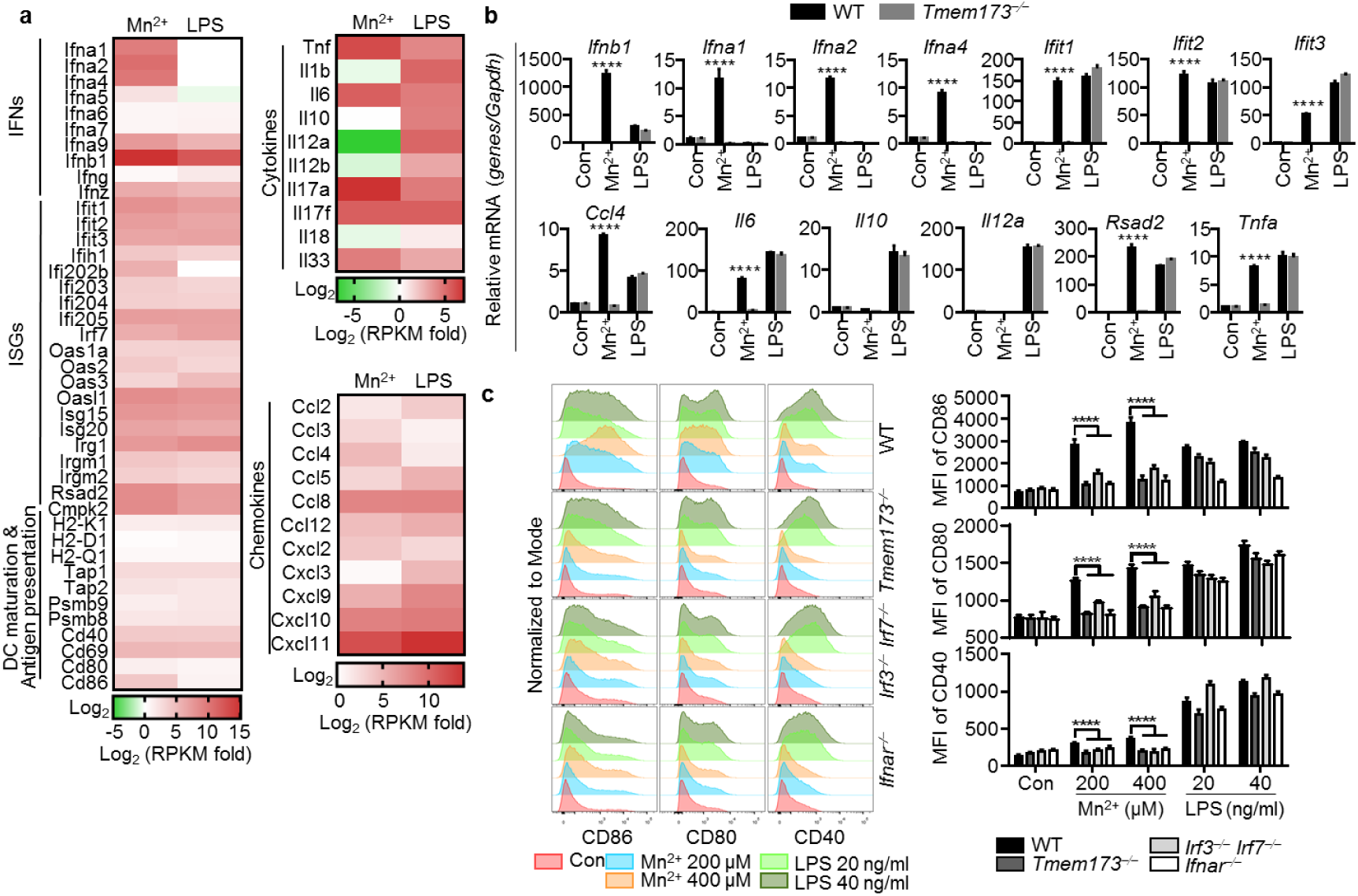
Mn^2+^ Promotes DC Maturation via cGAS-STING Activation. **a,** Heatmap of RNA-seq analysis. BMDCs were untreated or treated with MnCl_2_ (200 μM) or LPS (100 ng/ml) for 20 h. Heatmap was made by calculating log2 ((treated RPKM)/(control RPKM)). **b,** Quantitative RT-PCR analysis of the indicated gene expression in the WT and *Tmem173*^−⁄−^ BMDCs treated with MnCl_2_ (200 μM) or LPS (100 ng/ml) for 20 h. **c,** BMDCs from the WT, *Tmem173*^−⁄−^, *Irf3*^−⁄−^*Irf7*^−⁄−^ or *Ifnar*^−⁄−^ mice were treated with the indicated concentrations of MnCl_2_ or LPS for 20 h. CD86, CD80 and CD40 expression was analyzed by FACS. One representative experiment of at least three independent experiments is shown, and each was done in triplicate. Error bars represent SEM; **b,** data were analyzed by two-way ANOVA; **c,** data were analyzed by an unpaired t test. ns, not significant; * P < 0.05; ** P < 0.01; *** P < 0.001; **** P < 0.0001.

### Mn^2+^ Activates NLRP3 Inflammasome

Alum has been shown to activate the NLRP3 inflammasome and the capacity of Alum to promote antibody production was compromised in *Caspase1*^−⁄−^, *Pycard*^−⁄−^ or *Nlrp3*^−⁄−^ mice ^9–11^. To compare the ability of Mn^2+^ and Aluminium-containing adjuvants (Imject® Alum, Alhydrogel® adjuvant and Adju-Phos® adjuvant) to activate innate immunity, we tested the production of type I-IFNs and pro-inflammatory cytokines IL-1β and IL-18 in peritoneal macrophages treated with Mn^2+^ or various Alums. Only Mn^2+^ induced type I-IFNs (Fig. 2a). Mn^2+^ also activated stronger inflammasome than Imject® Alum and Alhydrogel® Alum did (Fig. 2b, c), which was entirely dependent on ASC, mainly on NLRP3 (Fig. 2d-g). Interestingly, Mn^2+^-induced inflammasome activation in mouse cells was different from THP1 cells, which showed a complete NLRP3-dependence (Fig. 2h, Extended Data Fig. 1a). Moreover, cGAS or STING deficiency did not affect Mn^2+^-activated inflammasome in macrophages from *Cgas*^−⁄−^, *Tmem173*^−⁄−^ mice or *CGAS*^−⁄−^, *TMEM173*^−⁄−^ THP1 cells (Extended Data Fig. 1b-e), indicating that cGAS-STING-induced lysosomal cell death ^29^, or cGAMP production ^30^ was not essential for Mn^2+^-induced inflammasome. Instead, we found that N-acetyl-L-cysteine (NAC, a direct scavenger of ROS), reduced L-glutathione (GSH, an intracellular thiol antioxidant), extracellular K^+^, or 2-APB (a cytosolic Ca^2+^ release inhibitor) all restrained Mn^2+^-induced inflammasome activation (Extended Data Fig. 2a-g). Using a modified culture medium, Hanks’ Balanced Salt Solution Deletion of PO_4_^3-^ and CO_3_^2-^ (herein HBSSD), in which Mn^2+^ and Ca^2+^ did not form particles, we found that Mn^2+^ activated NLRP3 and pyroptosis were essentially independent of particle formation, which is different from Ca^2+^ (Extended Data Fig. 2h, i). mtDNA depletion (Extended Data Fig. 2j) by ethidium bromide ^28^ did not affect Mn^2+^-induced inflammasome activation either (Extended Data Fig. 2k, l). We thus concluded that Mn^2+^ activated stronger NLRP3 inflammasome than Alum did.

**Fig. 2.**
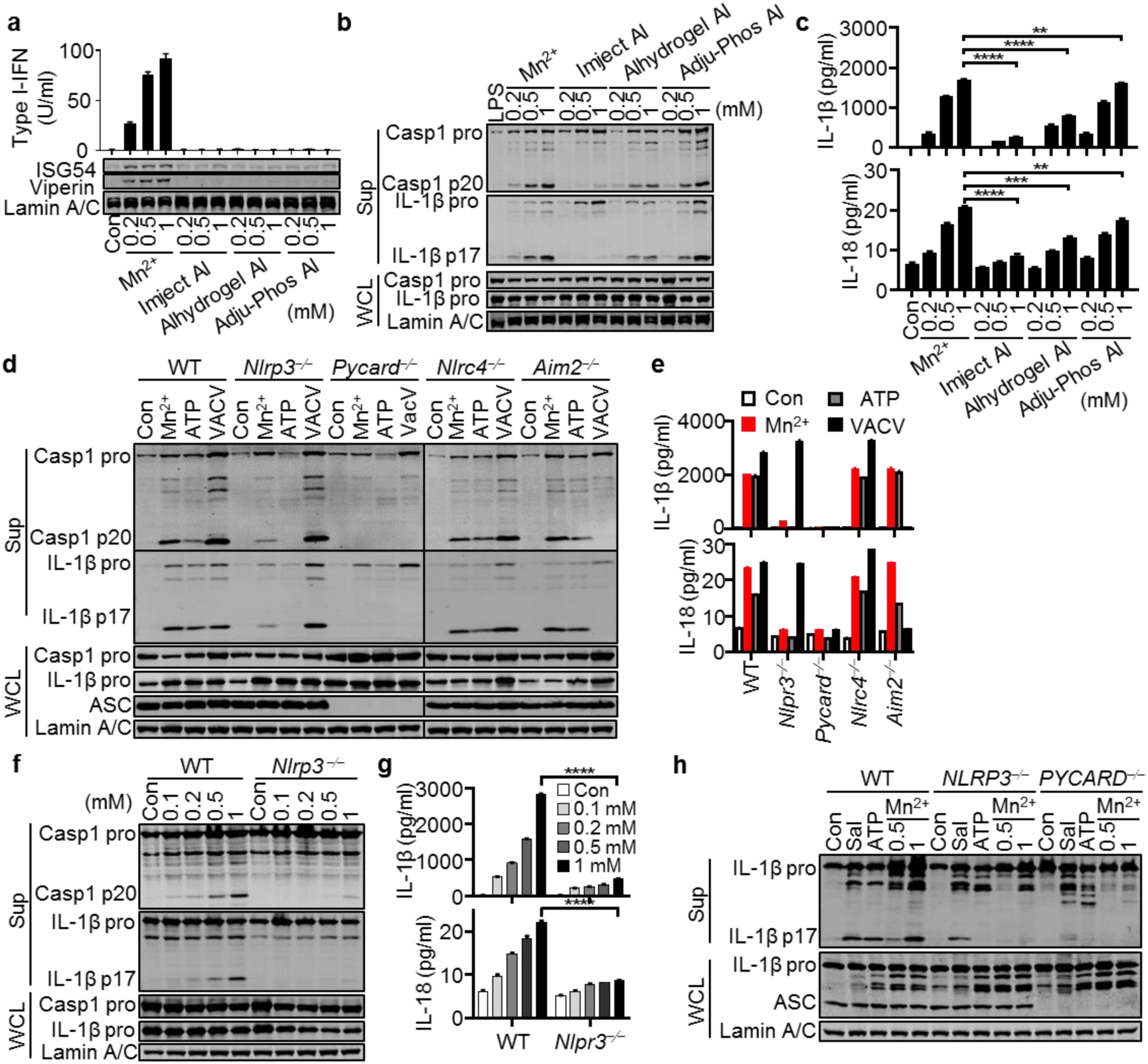
Mn^2+^ Activates NLRP3 Inflammasome. **a,** Western blot (lower) and Type I-IFN production analysis (upper) of mouse peritoneal macrophages treated with the indicated concentrations of MnCl_2_ or Aluminium salts (Imject® Alum, Alhydrogel® adjuvant 2%, Adju-Phos® adjuvant) for 18 h. **b, c,** Western blot (**b**) and ELISA analysis (**c**) of inflammasome activation of LPS-primed C57BL/6 peritoneal macrophages treated with the indicated concentrations of MnCl_2_ or Aluminium salts for 5 h. Supernatants (Sup) and whole cell lysates (WCL) were analyzed by immunoblotting with the indicated antibodies. **d, e,** Western blot (**d**) and ELISA analysis (**e**) of inflammasome activation of LPS-primed WT, *Nlrp3*^−⁄−^, *Pycard*^−⁄−^, *Nlrc4*^−⁄−^ and *Aim2*^−⁄−^ peritoneal macrophages treated with MnCl_2_ (0.5 mM), ATP (5 mM) or VACV (MOI = 10). **f, g,** Western blot (**f**) and ELISA analysis (**g**) of inflammasome activation of LPS-primed WT and *Nlrp3*^−⁄−^ peritoneal macrophages treated with the indicated concentrations of MnCl_2_ for 5 h. **h,** Western blot analysis of inflammasome activation of LPS-primed THP1 cells treated with Salmonella (Sal, MOI = 10), ATP (5 mM) or MnCl_2_ (0.5 and 1 mM). One representative experiment of at least three independent experiments is shown, and each was done in triplicate. Error bars represent SEM; **a, c, g,** data were analyzed by an unpaired t test. ns, not significant; * P < 0.05; ** P < 0.01; *** P < 0.001; **** P < 0.0001.

### Mn^2+^ Activates Inflammasome without IL-1/-18 Production

Consistent with results from RNA-seq and qPCR (Fig. 1a, b), Mn^2+^ treatment did not induce upregulation of *Il1b* and *Il18* in both murine BMDCs and human monocyte derived dendritic cells (Mo-DCs) (Fig. 3a). Interestingly, Alum activated NLRP3 inflammasome in a similar way, neither Mn^2+^ nor Alum induced IL-1/18 production without LPS priming (Fig. 3b, c). The same results were obtained when human peripheral blood mononuclear cells (PBMCs) were treated with Mn^2+^ (Fig. 3d). Since clinical studies on various inflammatory diseases suggested the crucial role for IL-1/-18 but not for TNFα, which only amplified and perpetuated the damage ^31^, we held that Mn^2+^-activated inflammasome did not cause systematic inflammation but may still be critical for its adjuvant activity (see below).

**Fig. 3.**
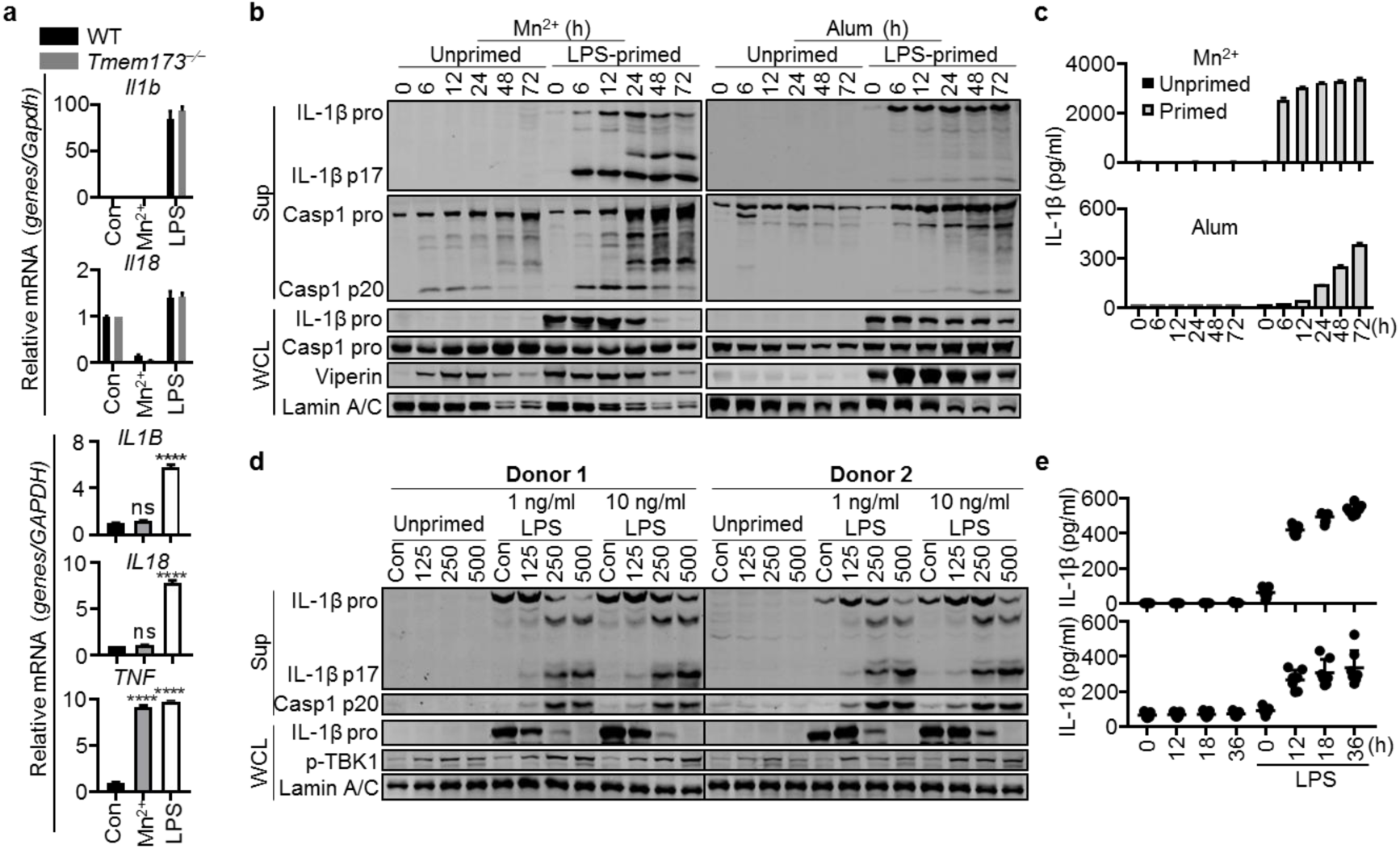
Mn^2+^ Activates Inflammasome without IL-1/-18 Production. **a,** Quantitative RT-PCR analysis of the indicated gene expression in the WT and *Tmem173*^−⁄−^ BMDCs (upper) or human Mo-DCs (lower) treated with MnCl_2_ (200 μM) or LPS (100 ng/ml) for 20 h. **b, c,** Unprimed or LPS-primed mouse peritoneal macrophages were treated with MnCl_2_ (200 μM) or Alum (100 μg/ml) for the indicated times. Supernatants (Sup) and whole cell lysates (WCL) were analyzed by immunoblotting with the indicated antibodies (**b**). IL-1β in supernatants was analyzed by ELISA (**c**). **d, e,** Unprimed or LPS (1 ng/ml or 10 ng/ml)-primed human PBMCs were treated with MnCl_2_ (200 μM) for the indicated times. Supernatants and whole cell lysates were analyzed by immunoblotting with the indicated antibodies (**d**). Human IL-1β and IL-18 in supernatants were analyzed by ELISA (**e**). One representative experiment of at least three independent experiments is shown, and each was done in triplicate. Error bars represent SEM; **a,** data were analyzed by an unpaired t test. ns, not significant; * P < 0.05; ** P < 0.01; *** P < 0.001; **** P < 0.0001.

### MnJ Is A Potent Universal Adjuvant

Given that Mn^2+^ induced strong type I-IFN production and NLRP3 inflammasome activation, we reasoned that Mn^2+^ could be used as an adjuvant. To test this, we first immunized C57BL/6 mice with LPS-free chicken ovalbumin protein (OVA) alone or OVA with different Mn^2+^ solutions intramuscularly (i.m.) or intranasally (i.n.) and measured OVA-specific antibodies. Surprisingly, we found that only Mn^2+^ in phosphate buffer saline (PBS), but not Mn^2+^ in normal saline, promoted antibody production (Extended Data Fig. 3a, b). The difference was that Mn^2+^ formed particles in PBS but not in saline, suggesting that soluble Mn^2+^ was unable to induce a local immune response as expected. However, Mn^2+^ particles in PBS tended to aggregate and precipitate with time, thus lost its adjuvant activity (Extended Data Fig. 3c). By screening various manganese compounds, we generated jelly-like Mn^2+^ colloids (MnJ, Mn Jelly) (Extended Data Fig. 3d) consisting of elongate nanoparticles approximately 2 × 2 × 6 nm in size (Extended Data Fig. 3e). MnJ was stable without aggregation in the following experiments. Its adjuvant effect was not weakened after storage at -80 °C for three weeks or longer, or even after freeze– drying/lyophilization (Extended Data Fig. 3f). On the contrary, Alum vaccines are known to be sensitive to freezing and thus hard to store or transport. Interestingly, MnJ triggered comparable inflammasome activation (Extended Data Fig. 3g), but stronger type I-IFN production than MnCl_2_ or Mn^2+^-PBS did (Extended Data Fig. 3h), probably due to the facilitated transportation into cells as nanoparticles ^32, 33^. Compared to MnCl_2_, which disappeared within hours, MnJ had a much longer muscle retention time up to 8 days at the site of injection, similar to Mn^2+^-PBS particles (Extended Data Fig. 3i). Splenocytes were isolated from OVA-immunized mice and stimulated with major histocompatibility class II (MHC-II)-binding OVA peptide I-A^b^ and MHC-I-binding peptide H-2K^b^ to compare T cell activation. IFNγ induced in splenocytes from OVA-MnJ was higher than OVA-Mn^2+^ (Extended Data Fig. 3j). Consequently, the adjuvant effect of MnJ was significantly better than that of Mn^2+^-PBS (Extended Data Fig. 3a-c).

We next evaluated the detailed adjuvant effect of MnJ. MnJ-adjuvanted antibody induction by intramuscularly immunization showed dose-dependence and lasted for at least 6 months (Fig. 4a). Compared to three different Aluminum-containing adjuvants, MnJ induced much stronger OVA-specific IgG1 production and CTL response (Fig. 4b). We also compared MnJ with other adjuvants including the complete Freund’s adjuvant (CFA), incomplete Freund’s adjuvant (IFA), MF59 and polyetherimide (PEI). We found that MnJ adjuvant effect was even better than CFA (20 μg MnJ vs 50 μl CFA, i.m.) and MF59 (20 μg MnJ vs 50 μl MF59, i.m.) (Extended Data Fig. 3k, l). Also, MnJ boosted specific antibody production against different recombinant protein/peptide antigens, including influenza A/PR8 hemagglutinin A1 peptide, haptenized experimental antigen nitrophenol-conjugatd keyhole limpet hemocyanin (NP-KLH), HBV surface antigen (HBsAg) and HBSS1 fusion protein (containing S (1-223 aa) and PreS1 (21-47 aa)) ^34^, and inactivated enterovirus type 71 (EV71) (Fig. 4d). Besides, intranasally immunization revealed that MnJ was also a potent mucosal adjuvant, inducing high levels of IgA antibodies in lung, saliva, and serum for a long time and as good as cholera toxin B (5 μg MnJ vs 10 μg CTB, i.n.) ^35^ (Fig. 4e, f).

**Fig. 4.**
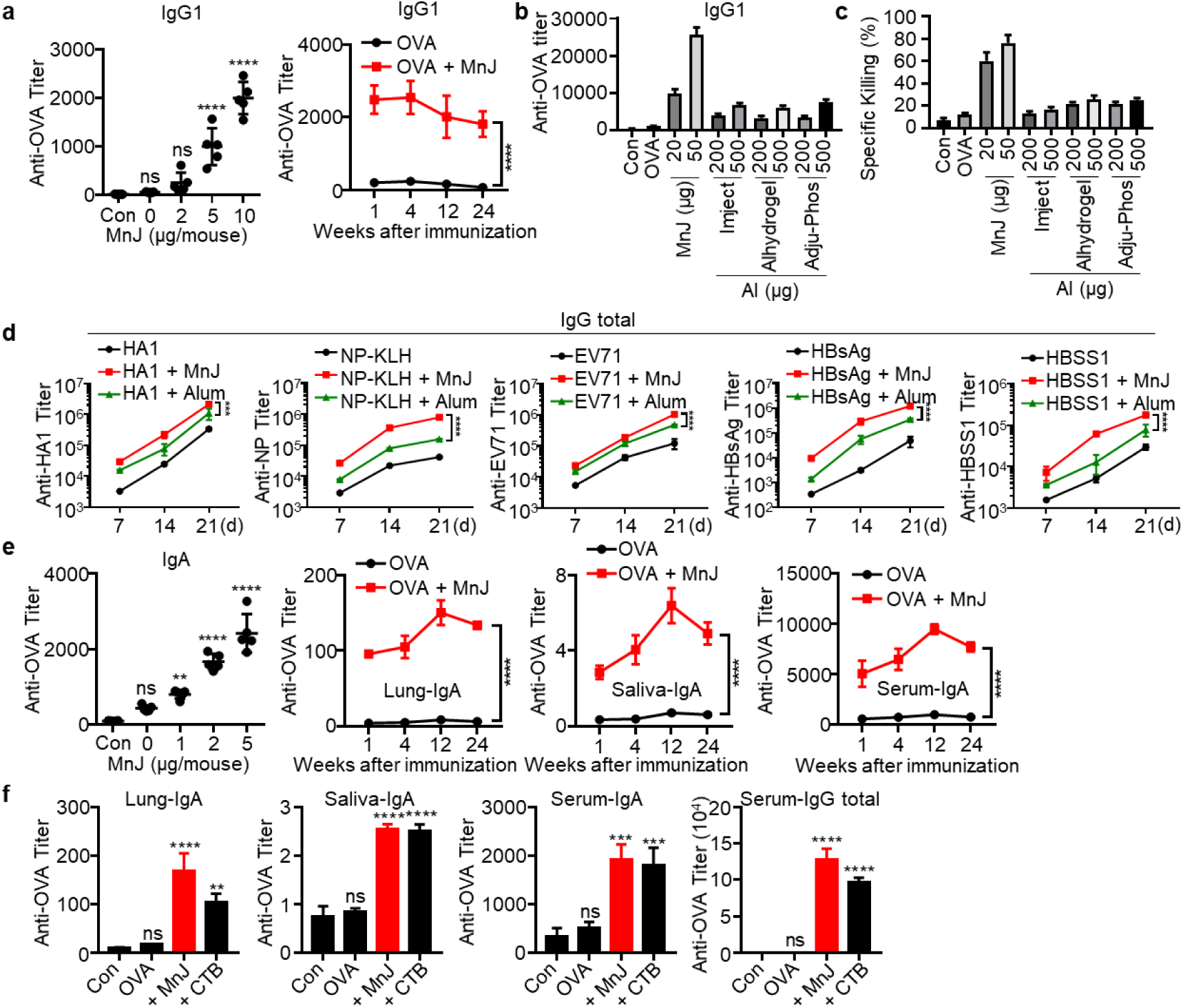
MnJ Is A Potent Universal Adjuvant. **a,** The WT mice (C57BL/6) were immunized intramuscularly with PBS or OVA (10 μg) + MnJ (0, 2, 5 or 10 μg) on day 0, 7 and 14. Sera were collected on day 21 to measure OVA-specific IgG1 by ELISA (left, n = 5). Time course of OVA-specific IgG1 in sera from mice immunized intramuscularly with OVA (10 μg) + MnJ (10 μg) for three times (right, n = 3). **b, c,** The WT mice were immunized intramuscularly with PBS, OVA (10 μg), OVA (10 μg) + MnJ (5 μg) or OVA (10 μg) + indicated amounts of Aluminium salts. Sera were collected on day 21 to measure OVA-specific IgG1 by ELISA (**b**) (n = 4). OVA-specific cytotoxicity was measured on day 21 in an *in vivo* killing assay (**c**) (immunized on day 0, 7 and 14, n = 4). **d,** The WT mice were immunized intramuscularly with the indicated antigen (5 μg), antigen (5 μg) + MnJ (10 μg) or antigen (5 μg) + Alum (1320 μg) on day 0, 7 and 14. Sera were collected on day 7, 14 and 21 to measure HA1, NP, EV71, HBsAg or HBSS1-specific IgG total (n = 3). **e,** The WT mice were immunized intranasally with PBS or OVA (10 μg) + MnJ (0, 1, 2 or 5 μg) on day 0, 7 and 14. Sera were collected on day 21 to measure OVA-specific IgA by ELISA (left, n = 5). Time course of OVA-specific IgA in BALF, saliva and serum as indicated from mice immunized intranasally with OVA (10 μg) + MnJ (5 μg) for three times (n = 3). **f,** OVA-specific IgA and IgG total were measured by ELISA on day 21 after immunization with OVA (10 μg), OVA (10 μg) + MnJ (5 μg) or OVA (10 μg) + CTB (10 μg) intranasally on day 0, 7 and 14 (n = 3). One representative experiment of at least three independent experiments is shown, and each was done in triplicate. Error bars represent SEM; **d,** data were analyzed by two-way ANOVA; **a, e, f,** data were analyzed by an unpaired t test. ns, not significant; * P < 0.05; ** P < 0.01; *** P < 0.001; **** P < 0.0001.

Importantly, compared to CFA-injected mice showing prominent swellings and granulomas with one shoot (Extended Data Fig. 4a), MnJ-injected mice displayed no visible side effects on injection site, body weight, survival or different organs even after repeated administrations (3 shoots in 3 consecutive weeks) (Extended Data Fig. 4a-i), suggesting that MnJ is a safe adjuvant with good biocompatibility.

### MnJ Promotes Antigen Presentation and T Cell Responses

MF59 and Alum facilitate APCs to engulf antigens and transport them to draining lymph nodes (dLNs), and also induce the differentiation of monocytes to dendritic cells (Mo-DCs) ^36, 37^. So, we next evaluated antigen uptake by APCs and Mo-DC differentiation in dLNs after MnJ administration. We immunized mice in both inguinal regions subcutaneously with fluorescent protein Phycoerythrin (PE) containing MnJ or Alum adjuvant. APCs in inguinal lymph nodes were analyzed by flow cytometry afterwards. The percentage and number of PE-loaded APCs were significantly enhanced 12 and 24 h after MnJ immunization (Fig. 5a). Also, there was a significant increase in the accumulation of Mo-DCs in mice immunized with antigen plus MnJ compared to that in mice immunized with antigen alone (Fig. 5b).

**Fig. 5.**
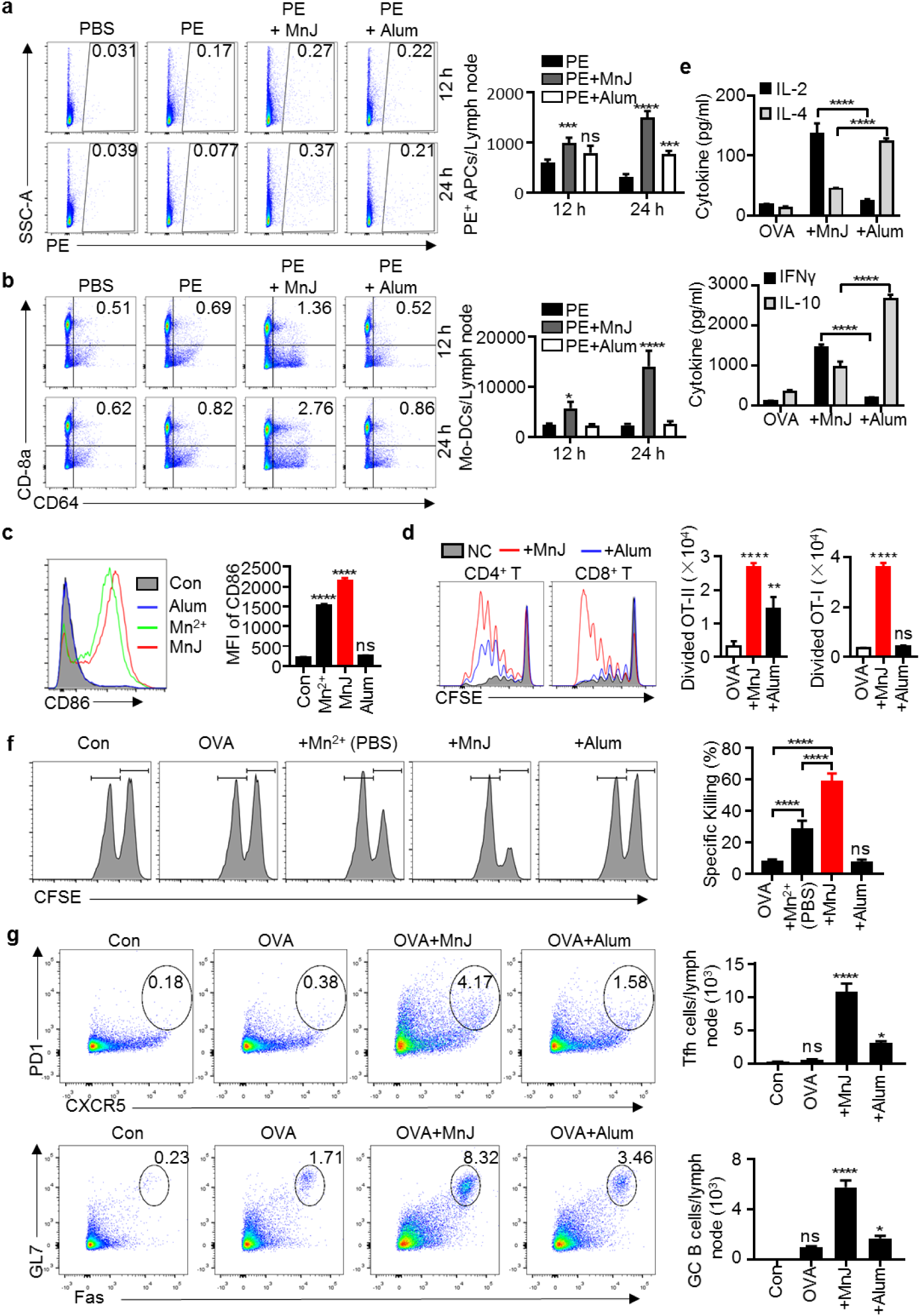
MnJ Promotes Antigen Presentation and T Cell Responses. **a, b,** The WT mice were immunized subcutaneously with PBS, Phycoerythrin (PE, 10 μg), PE (10 μg) + MnJ (20 μg) or PE (10 μg) + Alum (200 μg). Inguinal lymph nodes were collected 12 or 24 h later (n = 3). Ratio and number of PE^+^ APCs (**a**) and Mo-DCs (**b**) in dLN cells were analyzed by FACS. Live cells were identified by DAPI staining. Among live singlet cells, APCs were identified as the cell subset single or double positive for CD11c and F4/80. Among APCs, Mo-DCs were identified as CD8a^+^ CD64^+^ cells. **c,** BMDCs were treated with MnCl_2_ (20 μg/ml), MnJ (20 μg/ml) or Alum (20 μg/ml) for 20 h. CD86 expression was analyzed by FACS. **d,** CD45.1^+^ OT-I CD8^+^ or OT-II CD4^+^ T cells were labeled with CFSE and transferred to CD45.2^+^ WT mice. These mice were then immunized with OVA (1 μg), OVA (1 μg) + MnJ (10 μg) or OVA (1 μg) + Alum (1320 μg). After 3 days, T cell proliferation was analyzed by FACS (n = 3). **e,** The WT mice were immunized intramuscularly with OVA (10 μg), OVA (10 μg) + MnJ (10 μg) or OVA (10 μg) + Alum (1320 μg) on day 0, 7 and 14. Splenocytes were collected on day 21, and stimulated with OVA (100 μg/ml). IL-2, IL-4, IFNγ and IL-10 secreted by T cells were measured by ELISA (n = 3). **f,** OVA-specific cytotoxicity was measured on day 21 in an *in vivo* killing assay (immunized on day 0, 7 and 14, n = 4). **g,** Numbers of Tfh or GC B cells in dLN from WT mice were analyzed by FACS. Live cells were identified by DAPI staining. Among live singlet cells, CD4^+^ T cells were identified as the cell subset double positive for CD3 and CD4. Among CD4^+^ T cells, Tfh cells were identified as PD1^+^ CXCR5^+^ cells. B cells were identified as the cell subset double positive for CD45 and B220. Among B cells, GC B cells were identified as Fas^+^ GL7^+^ cells. One representative experiment of at least three independent experiments is shown, and each was done in triplicate. Error bars represent SEM; **a, b, e,** data were analyzed by two-way ANOVA; **c, d, f, g,** data were analyzed by an unpaired t test. ns, not significant; * P < 0.05; ** P < 0.01; *** P < 0.001; **** P < 0.0001.

The capacity of MnJ to promote BMDC maturation was stronger compared with Mn^2+^, while Alum did not have any effect (Fig. 5c). *In vivo*, MnJ enhanced both CD4^+^ and CD8^+^ T cell proliferation, whereas Alum only induced a weak CD4^+^ T cell proliferation (Fig. 5d). Splenocytes were next isolated from OVA-immunized mice and stimulated with OVA to compare T cell activation. IL-2 and IFNγ were highly induced in splenocytes from OVA-MnJ, but not OVA-Alum immunized mice, whereas IL-4 and IL-10 were preferably produced via OVA-Alum immunization (Fig. 5e), indicating that MnJ potently stimulated TH1 response, in addition to TH2 response. Accordingly, *in vivo* cytotoxic assay showed that MnJ immunized mice generated very strong CTL activities in killing OVA-bearing cells, which was absent in Alum immunized mice (Fig. 5f). MnJ also promoted the formation of germinal center (GC) with significantly increased amounts of Tfh and GC B cells (Fig. 5g).

### Both cGAS-STING and NLRP3 Inflammasome Contribute to Adjuvant Activity of MnJ

Next, *Tmem173*^−⁄−^, *Mavs*^−⁄−^, *Nlrp3*^−⁄−^, *Nlrc4*^−⁄−^, *Aim2*^−⁄−^ or *Pycard*^−⁄−^ mice were used to test which pathway is involved in MnJ’s adjuvant effect. It was found that although *Tmem173*^−⁄−^, *Pycard*^−⁄−^ or *Nlrp3*^−⁄−^ mice produced diminished OVA-specific antibodies (Extended Data Fig. 5a), *Tmem173*^−⁄−^*Pycard*^−⁄−^ (double knockout, DKO) mice generated extremely decreased OVA-specific antibodies (Fig. 6a-c), suggesting that both cGAS-STING and inflammasome contributed to MnJ adjuvant effect. Further, *Tmem173*^−⁄−^*Nlrp3*^−⁄−^ mice generated a bit more antibodies than *Tmem173*^−⁄−^*Pycard*^−⁄−^ mice did (Fig. 6a-b), indicating the involvement of other ASC-dependent inflammasome activation by MnJ. We also analyzed the expression of *Dock2*, which was reported to be down-regulated in *Pycard*^−⁄−^ but not *Nlrp3*^−⁄−^ or *Caspase1*^−⁄−^ mice ^38^. Quantitative PCR analysis confirmed the same expression of *Dock2* mRNA in WT, *Tmem173*^−⁄−^, *Pycard*^−⁄−^, *Nlrp3*^−⁄−^, *Tmem173*^−⁄−^*Pycard*^−⁄−^ and *Tmem173*^−⁄−^*Nlrp3*^−⁄−^ mice (Extended Data Fig. 5b). Consistently, MnJ-promoted germinal center formation was impaired in the DKO (specifically referred to *Tmem173*^−⁄−^*Pycard*^−⁄−^ in the following text) mice (Extended Data Fig. 5c). In addition, MnJ-induced OT-II CD4^+^ T cell proliferation disappeared in the DKO mice (Fig. 6d), along with sharply reduced IFNγ and IL-2 production by peptide-stimulated splenic T cells (Fig. 6e). Consistent with previous results (Fig. 3d, e), no IL-1β induction was detected in lymph nodes from MnJ-treated mice *in vivo*, despite of ISG production and GSDMD cleavage (Fig. 6f). Since neither Mn^2+^ nor MnJ treatment did induce IL-1 or IL-18 expression *in vitro* or *in vivo*, we reasoned that ASC-mediated inflammasome activation contributed to MnJ adjuvant activity by releasing other DAMPs like uric acid ^39^, but not by IL-1 or TNFα, consistent with previous reports that neither of them was important for adjuvant effect ^22, 40^. However, CFA or Monophosphoryl Lipid A (MPLA) induced antibody production did not change much among these mice (Extended Data Fig. 5d, e).

**Fig. 6.**
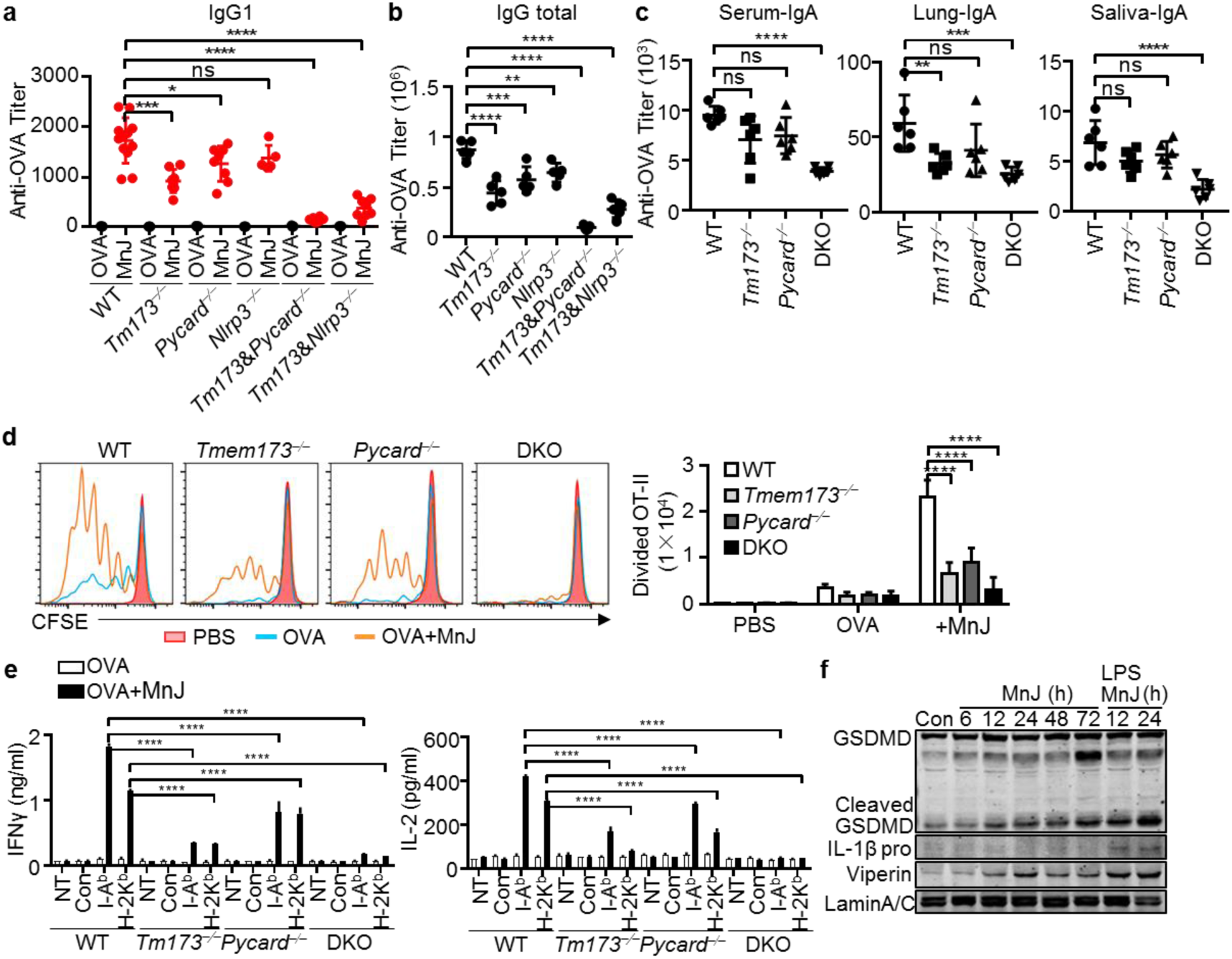
Both cGAS-STING and NLRP3 Inflammasome Contribute to Adjuvant Activity of MnJ. **a, b,** OVA-specific IgG1 (**a**) and IgG total (**b**) from the WT, *Tmem173*^−⁄−^, *Pycard*^−⁄−^, *Nlrp3*^−⁄−^, *Tmem173*^−⁄−^*Pycard*^−⁄−^ and *Tmem173*^−⁄−^*Nlrp3*^−⁄−^ mice were measured by ELISA on day 14 after immunization with OVA (10 μg) or OVA (10 μg) + MnJ (10 μg) intramuscularly on day 0 and 7 (n > 5). **c,** OVA-specific IgA from the WT, *Tmem173*^−⁄−^, *Pycard*^−⁄−^ and *Tmem173*^−⁄−^*Pycard*^−⁄−^ DKO mice was measured by ELISA on day 21 after immunization with OVA (10 μg) or OVA (10 μg) + MnJ (5 μg) intranasally on day 0, 7 and 14 (n = 6). **d,** CD45.1^+^ OT-II CD4^+^ T cells were labeled with CFSE and transferred to CD45.2^+^ WT, *Tmem173*^−⁄−^, *Pycard*^−⁄−^ and *Tmem173*^−⁄−^*Pycard*^−⁄−^ DKO mice. These mice were then immunized with PBS, OVA (1 μg) or OVA (1 μg) + MnJ (10 μg). After 3 days, T cell proliferation was analyzed by FACS (n = 3). **e,** The WT, *Tmem173*^−⁄−^, *Pycard*^−⁄−^ and *Tmem173*^−⁄−^*Pycard*^−⁄−^ DKO mice were immunized as (**a**). Splenocytes were collected on day 21, and stimulated with OVA peptides. IFNγ and IL-2 secreted by T cells were measured by ELISA (n = 3). **f,** The WT mice were immunized with MnJ (100 μg) or LPS (20 μg) + MnJ (100 μg) intramuscularly for the indicated times. Lysates of draining lymph node cells were analyzed by immunoblotting with the indicated antibodies. One representative experiment of at least three independent experiments is shown, and each was done in triplicate. Error bars represent SEM; data were analyzed by an unpaired t test. ns, not significant; * P < 0.05; ** P < 0.01; *** P<0.001; **** P < 0.0001.

### MnJ Is A Potent Adjuvant for Antiviral and Antitumor Vaccines

Next we tested the protection effect of MnJ adjuvanted vaccines against various viruses. Formaldehyde-inactivated Vesicular stomatitis virus (VSV), Herpes simplex virus 1 (HSV-1) and Vaccinia Virus (VACV) were used as vaccines. The optimal dose for each inactivated virus was first determined (VSV and HSV i.m.; VACV i.n.) (Extended Data Fig. 6a-f). MnJ greatly enhanced the protection efficacy of inactivated virus by about 100 times, which is much better than Alum. Consistently, virus titers were decreased by at least 10^3^ times in MnJ immunized mice (Fig. 7a-f), indicating that MnJ can be applied to all tested virus vaccines through either intramuscular or intranasal immunization, with greatly reduced amounts of inactivated-virus needed for an adequate protection.

**Fig. 7.**
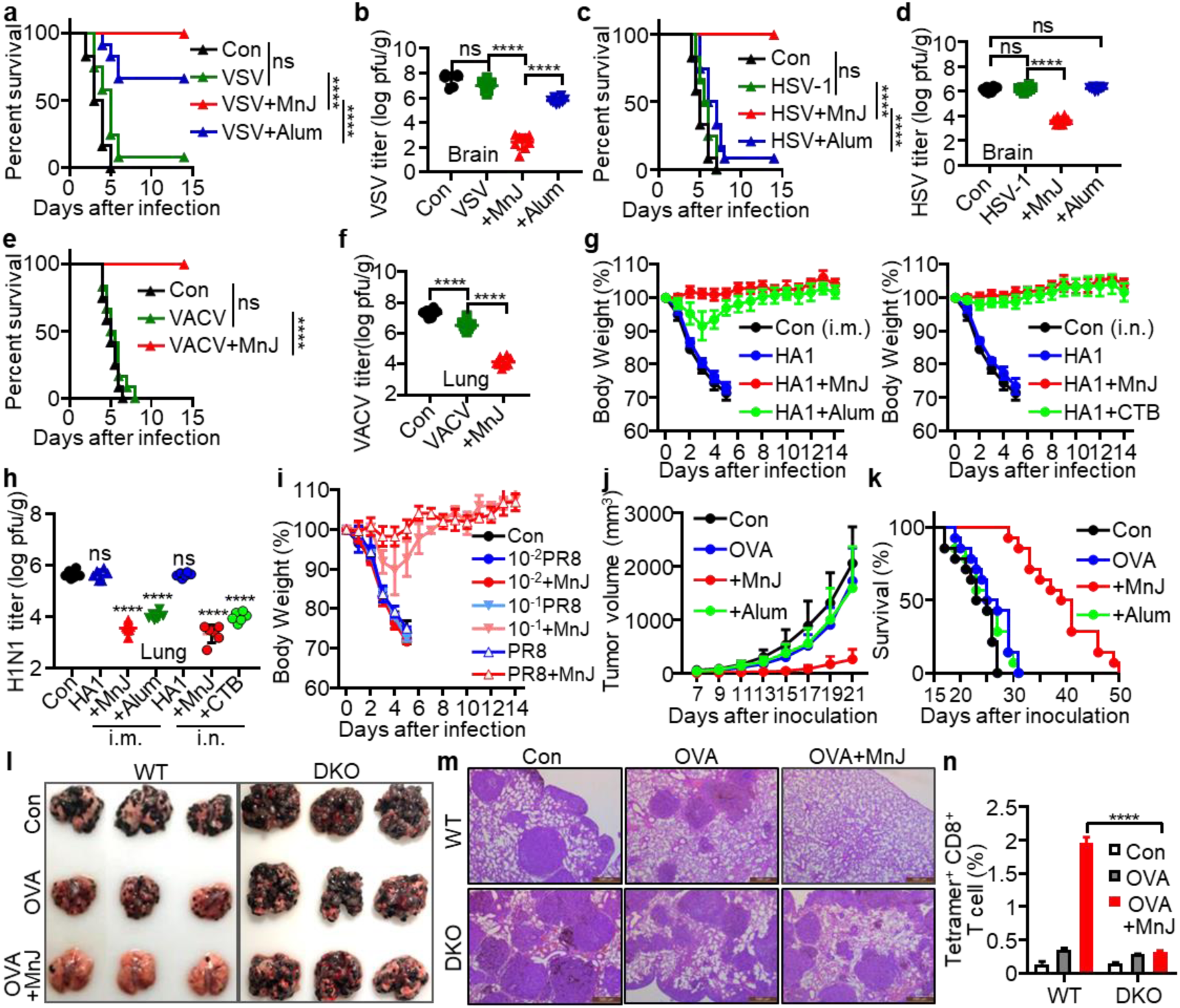
MnJ Is A Potent Adjuvant for Antiviral and Antitumor Vaccines. **a, b,** The WT mice were immunized intramuscularly with PBS, inactivated VSV (10^5^ pfu), inactivated VSV (10^5^ pfu) + MnJ (10 μg) or inactivated VSV (10^5^ pfu) + Alum (1320 μg) on day 0. On day 10, these mice were infected intravenously with a lethal dose of VSV. The survival was monitored for 2 weeks (n = 12) (**a**). Viral loads in brain were measured 5 days after infection (n = 8) (**b**). **c, d,** The WT mice were immunized intramuscularly with PBS, inactivated HSV-1 (10^3^ pfu), inactivated HSV-1 (10^3^ pfu) + MnJ (10 μg) or inactivated HSV-1 (10^3^ pfu) + Alum (1320 μg) on day 0. On day 10, these mice were infected intraperitoneally with a lethal dose of HSV-1. The survival was monitored for 2 weeks (n = 12) (**c**). Viral loads in brain were measured 5 days after infection (n = 8) (**d**). **e, f,** The WT mice were immunized intranasally with PBS, inactivated VACV (2 × 10 ^4^ pfu) or inactivated VACV (2 × 10^4^ pfu) + MnJ (5 μg) on day 0 and 7. On day 14, these mice were infected intranasally with a lethal dose of VACV. The survival was monitored for 2 weeks (n = 12) (**e**). Viral loads in lung were measured 5 days after infection (n = 8) (**f**). **g,** The WT mice were immunized intramuscularly (i.m.) with HA1 (5 μg), HA1 (5 μg) + MnJ (10 μg), HA1 (5 μg) + Alum (1320 μg) or intranasally (i.n.) with HA1 (5 μg), HA1 (5 μg) + MnJ (10 μg), or HA1 (5 μg) + CTB (10 μg) on day 0, 7 and 14. On day 21, these mice were infected intranasally with a lethal dose of PR8. The survival was monitored for 2 weeks (n = 10). **h,** The WT mice were immunized intranasally with PBS, inactivated PR8 (5 × 10^6^ pfu), 10^-1^ PR8 (5 × 10^5^ pfu) or 10^-2^ PR8 (5 × 10^4^ pfu) with or without MnJ (5 μg) on day 0 and 7. On day 14, these mice were infected intranasally with a lethal dose of PR8. The body weight was recorded for 2 weeks (n = 3). **i,** Viral loads in lungs from mice in (**g**) were measured 5 days after infection (n = 6). **j, k,** The WT mice were immunized intramuscularly with PBS, OVA (10 μg), OVA (10 μg) + MnJ (20 μg) or OVA (10 μg) + Alum (1320 μg) on day 0, 7 and 14. On day 21, these mice were inoculated with B16-OVA-Fluc cells (3 × 10^5^) subcutaneously. Tumor volume (**j**) was measured and survival (**k**) was monitored (n = 14). **l, m, n,** The WT and DKO mice were immunized intramuscularly with PBS, OVA (10 μg), or OVA (10 μg) + MnJ (20 μg) on day 0, 7 and 14. On day 21, these mice were inoculated with B16-F10-OVA (3 × 10^5^) intravenously. Images (**l**) and HE staining (**m**) of lung tissues were recorded 20 days after inoculation (n = 6). The percentage of tetramer^+^ CD8^+^ T cells in spleens of these mice was analyzed by FACS on day 21 (n = 3) (**n**). One representative experiment of at least three independent experiments is shown, and each was done in triplicate. Error bars represent SEM; **b, d, f, h, n,** data were analyzed by an unpaired t test; **a, c, e, k,** survival plot data were analyzed with log-rank (Mantel–Cox) tests. ns, not significant; * P < 0.05; ** P < 0.01; *** P<0.001; **** P < 0.0001.

We particularly tested MnJ in influenza vaccines. Mice immunized with virus protein HA1 (PR8) plus MnJ (i.m. or i.n.) or HA1 plus CTB (i.n.) were completely protected from lethal H1N1 A/PR8/34 virus challenge, while Alum adjuvant only showed mild protection (Fig. 7g, h, Extended Data Fig. 6g-i). Importantly, MnJ enhanced the protection effect of inactivated PR8 vaccine by at least 10 times (Fig. 7i). Moreover, MnJ exhibited superior protection against heterologous influenza viruses to mice immunized with MnJ-adjuvanted inactivated PR8 or HA1 protein, followed by lethal H1N1 WSN or H3N2 challenge (Extended Data Fig. 6j).

Finally, the adjuvant effect of MnJ on cancer vaccines was evaluated. Mice were immunized three times before inoculated with melanoma cell B16-OVA subcutaneously. Tumor growth was greatly suppressed in OVA-MnJ, but not OVA-Alum, immunized mice (Fig. 7j, Extended Data Fig. 7a), in line with prominently improved survival (Fig 7k) and increased tumor-infiltrating CD4^+^ and CD8^+^ T cells (Extended Data Fig. 7b, c), confirming CTL inducing activity of MnJ. In pulmonary metastasis model, OVA-MnJ immunization greatly blocked lung metastases in the WT, but not *Tmem173*^−⁄−^*Pycard*^−⁄−^ mice (Fig. 7l, m, Extended Data Fig. 7d), which is consistent with the proliferation of CD8^+^ OT-I T cells (Extended Data Fig. 7e). In addition, OVA-specific antibody production and CD8^+^ T cell activation analyzed by tetramer assays were only detected in MnJ-OVA immunized WT mice (Fig. 7n, Extended Data Fig. 7f, g), suggesting its potentials in tumor therapies.

## DISCUSSION

The adjuvant activity of Aluminum was found by Alexander Glenny and his colleagues in 1926 ^41^, now almost one century later, we reported the adjuvant activity of another metal-Manganese. Adjuvant effect is essentially attributed by type I-IFN-promoted dendritic cell maturation and migration to prime adaptive immune responses along with 1) local up-regulation of chemokines, including CCL2 (MCP-1) and CCL3 (MIP-1α) to recruit immune cells to the injection site; 2) increase of antigen uptake by immune cells; 3) induction of monocyte differentiation into dendritic cells ^8, 36, 37^. We found that Mn^2+^ induced APCs to produce both IFNβ and various IFNαs, which were surprisingly not induced by LPS, together with many co-stimulatory and MHC molecules crucial for antigen presentation and chemokines for immune cell recruitment. MnJ also strongly enhanced antigen uptake by APCs and the differentiation of monocytes to dendritic cells. Importantly, Mn^2+^ did not induce the production of pro-inflammatory cytokines IL-1 and IL-18 in human or mouse *in vitro* and *in vivo*. Therefore, MnJ demonstrated superior adjuvant effects as it induces humoral, cellular, and mucosal immune responses, particularly CTL activation, without detected side-effects. In addition, the adjuvant effect of MnJ was stable even after repeated freeze-drying cycles, whereas Alum is sensitive to freezing and requires cold-chain temperature control ^42, 43^. MnJ showed great adjuvant effects to all tested vaccine antigens including inactivated viruses, recombinant protein subunits and peptides, thus it can significantly reduce the amount of viruses needed. Particularly, its tumor antigen-specific CTL activity by either intramuscular or intranasal MnJ immunization indicated a great potential for cancer vaccines. Based on these results, we believe that MnJ has great potentials for the development of potent but safe vaccines. The component simplicity and steadiness of MnJ, the low cost and wide availability of Mn make this adjuvant even more promising.

Although Alum adjuvanted vaccines have been proven to be safe in most cases, some studies showed that aluminum accumulation are associated with long-lasting macrophagic myofasciitis (MMF) ^44^, nervous disoders ^45^ and bone disease ^46^ in some patients. The FDA-approved doses of 850 μg, 1140 μg, and 1250 μg Alum per vaccine were determined according to antigenicity and effectiveness of vaccine, not including safety consideration ^47–49^. After vaccine injection, Alum has been found in the injected muscle, draining lymph nodes and spleen 9 months in humans ^50^ or even 12 years in patients with ASIA (Autoimmune/inflammatory syndrome induced by adjuvants) ^51, 52^. Consistently, we found that intramuscularly injected Alum in mice did not show obvious clearance 2 weeks after injection whereas injected MnJ was beyond detection after 8 days. Importantly, MnJ, but not Alum, could be readily decomposed into free Mn^2+^ ions by many common acidic metabolic products or under acidic environment (data not shown). In addition, tumor microenvironment is characterized by anomalous metabolic properties and acidic environment ^53^, which might be beneficial for the application of MnJ in antitumor therapy. Excessive Mn accumulation in the central nervous system causes neurologic toxicity in occupational cohorts through inhalation from welding or smelting ^26^. However, we found that 20 μg MnJ induced higher antibody production than 1444 μg Imject® Alum, 1444 μg Alhydrogel® adjuvant 2%, 2259μg Adju-Phos® adjuvant (500 μg Alum in each) did, indicating MnJ is at least 25 times more potent than Alum in terms of inducing humoral immune response, which means that much smaller amount of MnJ can achieve the same effect.

There is still controversy about the function of NLRP3 inflammasome on adjuvant effect of aluminum salt ^12^. These different conclusions may result from different aluminum salts or mice backgrounds used. There is also another view that ASC has an inflammasome-independent role in shaping adaptive immunity, for regulating the expression of Dock2, which is important for antigen uptake and lymphocyte mobility ^38^. However, we found that STING, NLRP3 or ASC deficiency did not affect the expression of Dock2 in macrophages or lymphocytes. Interestingly, manganese salts administration did not upregulate pro-IL-1β or pro-IL-18 production, which means that MnJ activate adaptive immune responses partly through an inflammasome-dependent but inflammatory cytokines-independent manner. In this regard, there may be some other stimulators upregulated by MnJ and released by pyroptotic death of cells. However, even though we did not detect IL-1/-18 production by macrophages or PBMCs treated with MnCl_2_ alone, these cells may generate these cytokines when treated with inactivated pathogens plus MnCl_2_. IL-1 and IL-18 can promote the infiltration of neutrophils and enhance immune responses after immunization with adjuvants like Alum or ISCOMATRIX ^40, 54, 55^.

In addition, similar to Alum adjuvant, the MnJ nanoparticles were also able to absorbe antigens like OVA, GFP or PE proteins (data not shown). The physical depot effect of MnJ retained antigens at the injection site and enhanced the uptake of antigens by APCs. Generally, MnJ is an adjuvant owning the properties of both immune potentiator and delivery system. Because of its excellent adjuvant activities and stability against repeated freezing-thaw treatment, Mn-based adjuvants would be especially useful in veterinary vaccines with the following three additional advantages. 1) MnJ showed a very nice dose-dependent adjuvant activity intramuscularly or intranasally, with 20 μg MnJ per mouse showing stronger antibody inducing activity than any tested adjuvants including CFA; 2) High MnJ dose (up to several mg/shot) and repeated administrations to mice, rabbits or pigs (data not shown) did not cause any visible damage or inflammation such as swellings and granulomas at the injection sites or in various organs; 3) Mammals keep tissue Mn levels via tight control of both absorption and excretion, as normally only 1–5% of ingested Mn is absorbed into the body ^56^ and excessive dietary Mn causes reduced Mn absorption and enhanced Mn metabolism and excretion ^57–59^. Therefore, even highly elevated Mn levels in meats caused by Mn-containing veterinary vaccines would unlikely increase gastrointestinal Mn absorption by humans. In fact, Mn contents in whole grains, rice, and nuts are around or more than 30 mg Mn/kg or even 110-140 mg Mn/kg in wheat bran, much higher than those in mammals that are between 0.3 and 2.9 mg Mn/kg wet tissue weight, confirming the tight Mn absorption by animals.

## Acknowledgements

We thank Ms. Liying Du, Drs Hongxia Lv, Guilan Li, Xiaochen Li from National Center for Protein Sciences at Peking University in Beijing, China, for assistance with FACS, protein purification and microscopy. We thank Drs. Yan Shi for OT-I and OT-II mice, Zhijian Chen for *Mavs*^−⁄−^ mice, Rongbin Zhou for *Ifnar*^−⁄−^ mice, Tadatsugu Taniguchi for *Irf3*^−⁄−^ *Irf7*^−⁄−^ mice, Vishva M. Dixit for *Aim2*^−⁄−^, *Pycard*^−⁄−^, *Nlrc4*^−⁄−^ and *Nlrp3*^−⁄−^ mice, Hongbing Shu, Yonghui Zhang, Wenjun Liu, Min Fang for viruses, Yonghui Zhang, Wenhui Li, Wenjie Tan, Changfa Fan for antigens. This work was supported by National Natural Science Foundation of China (31830022 and 81621001) and the Chinese Ministry of Science and Technology (2015CB943203).

## Author contributions

R.Z., C.W. and Z.J. designed research; R.Z., C.W. and Y.G. performed the experiments, X.W., M.J., M.S., M.L., J.X. and Y.W. assisted in the experiments. R.Z., C.W. and Z.J. analyzed the data and wrote the manuscript.

## Competing interests

The authors declare no competing interests.

## Data and materials availability

All data supporting the findings of this study are available within the paper and its supplementary materials.

RNA-seq data have been deposited in Gene Expression Omnibus under accession no. GSE126586.

