## Supplementary material for "The Manganese Salt (MnJ) Functions as A Potent Universal Adjuvant": Supplymentary information

**This file includes:**

Materials and Methods

Figs. S1 to S7

Tables S1 to S3

References

### Materials and Methods

#### Human subjects

The study was approved by the Ethical Committee on Human Research of Peking University and was in accordance with the Declaration of Helsinki. PBMCs were isolated from peripheral blood of volunteers (Table S1) using Histopaque-1077 (Sigma, 10771) through consecutive centrifugation. PBMCs were seeded in 12-well plates at a final concentration of $5 \times 10^6$ cells/well with Opti-MEM (Gibco) and treated with the indicated concentrations of LPS or $Mn^{2+}$.

#### Mice

Wild-type (WT) C57BL/6 Specific Pathogen Free (SPF) mice were purchased from Vital River, China. *Cgas*$^{-/-}$ and *Tmem173*$^{-/-}$ mice have been previously described [1]. OT-I (C57BL/6-Tg (TcraTcrb) 1100Mjb/J) mice and OT-II (B6.Cg-Tg (TcraTcrb) 425Cbn/J) mice were gifts from Dr. Yan Shi. *Mavs*$^{-/-}$ mice were gifts from Dr. Zhijian Chen. *Irf3*$^{-/-}$*Irf7*$^{-/-}$ mice were gifts from Dr. Tadatsugu Taniguchi. *Ifnar*$^{-/-}$ mice were gifts from Dr. Rongbin Zhou. *Pycard*$^{-/-}$ mice, *Aim2*$^{-/-}$ mice, *Nlrp3*$^{-/-}$ mice and *Nlrc4*$^{-/-}$ mice were gifts from Dr. Vishva M. Dixit. *Tmem173*$^{-/-}$*Pycard*$^{-/-}$ DKO mice were generated by crossing *Tmem173*$^{-/-}$ mice and *Pycard*$^{-/-}$ mice.

All mice were bred and kept under specific pathogen-free conditions in the Laboratory Animal Center of Peking University in accordance with the National Institute of Health Guide for Care and Use of Laboratory Animals.

#### Cell lines

L929-ISRE, MDCK, BHK21, B16-F10-OVA, B16-F10-OVA-GFP cells were cultured in DMEM (Gibco) supplemented with 10% FBS (Gibco), 5 μg/ml of penicillin and 10 μg/ml of streptomycin. THP1 and its derived knockout cells were cultured in RPMI-1640 (Gibco) media supplemented with 10% FBS (Gibco). THP1 gene-specific knockout cells were generated by CRISPR-cas9 system with gRNA sequence in Table S2.

Peritoneal macrophages were harvested from mice 6 days after thioglycollate (BD, Sparks, MD) injection, and cultured in DMEM supplemented with 5% FBS.

For induction of immature BMDCs, bone marrow cells were isolated from WT and indicated knockout mice and cultured in RPMI-1640 (Gibco) media supplemented with 10% FBS (Gibco), 20 ng/ml of mIL-4 (Genscript) and 20 ng/ml of mGM-CSF (Genscript) at 37 ℃ and 5% $CO_2$. On day 3, half of the culture supernatant was changed to fresh medium. On day 7, non-adherent cells were collected and used immediately. DC purity was confirmed with PE labelled anti-CD11c (Biolegend) by FACS.

For induction of human Mo-DCs, monocytes were isolated from human PBMCs with MojoSort™ Human Pan Monocyte Isolation Kit (Biolegend) and cultured in RPMI-1640 (Gibco) media supplemented with 10% FBS (Gibco), 100 ng/ml of hIL-4 (Genscript) and 100 ng/ml of hGM-CSF (Genscript) at 37 ℃ and 5% $CO_2$. On day 3, half of the culture supernatant was changed to fresh medium. On day 7, non-adherent cells were collected and used immediately. DC purity was confirmed with APC labelled anti-CD11c (Biolegend) by FACS.

**Antibodies**

cDNA of Viperin (mouse 243 - 360 aa), ISG54 (mouse 3 - 289 aa), Casp1/p20 (mouse 121 - 296 aa), IL-1β/p17 (mouse 118 - 269 aa; human 117 - 269 aa), ASC (mouse full-length), cGAS (human full length), STING (human 140 - 379 aa), GSDMD (mouse 276 – 487 aa) and Lamin A/C (human full length) were cloned into pET-21b vector (Novagen) and proteins were expressed in *E. coli* BL21 (DE3). The recombinant protein was purified by Ni-NTA affinity chromatography, and then injected into mice or rabbits to produce antiserum. Anti-GAPDH (sc-25778) antibody was from Santa Cruz Biotechnology. Anti-cGAS (#31659) antibody was from Cell Signaling Technology. HRP-rat anti-mouse IgG1 polyclonal antibody (eBioscience, 18-4015-82), HRP-goat anti-mouse IgG2b polyclonal antibody (Southern Biotech, 1091-05), HRP-goat anti-mouse IgG2c polyclonal antibody (GeneTex, GTX77297), HRP-goat anti-mouse IgG3 polyclonal antibody (Southern Biotech, 1101-05), HRP-goat anti-mouse IgG (H+L) (Proteintech, SA00001), HRP-goat anti-mouse IgA polyclonal antibody (GeneTex, GTX77223) was purchased as indicated.

**Reagents**

Mouse IL-1β ELISA Kit (MULTI SCIENCES, 70-EK201B2/2), Mouse IL-18 ELISA Kit (MULTI SCIENCES, 70-EK2182), Human IL-1β ELISA Kit (Invitrogen, 88-7261-22), Human IL-18 ELISA Kit (Sino Biological, SEK 10119), Mouse IFNγ ELISA Kit (MULTI SCIENCES, 70-EK280/3-96), Mouse IL-2 ELISA Kit (MULTI SCIENCES, 70-EK2022/2), LPS (Sigma, L4130), Mouse IL-4 Uncoated ELISA Kit (Invitrogen, 88-7044-22), Mouse IL-10 Uncoated ELISA Kit (Invitrogen, 88-7105-22), Z-VAD-FMK (Selleck, S7023), ABT-263 (Selleck, S1001), Ovalbumin (InvivoGen, #vac-pova), OVA peptides specific for I-$A^b$ (ISQAVHAAHAEINEAGR) or H-$2K^b$ (SIINFEKL) or control peptide (FAPGNYPAL) were synthesized by Scilight Biotechnology, N-acetyl-L-cysteine (Sigma, A7250), Reduced L-glutathione (Sigma, G4251), 2-Aminoethyl diphenylborinate (Sigma, D9754), Influenza A H1N1 (A/PR8/34) Hemagglutinin / HA1 peptide (11-324 aa) and MF59 adjuvant were from Yonghui Zhang, Tsinghua University. HBsAg (small envelope protein, 226 aa) was from Wenhui Li, National Institute of Biological Sciences. EV71 inactivated virus (H07 strain) was from Changfa Fan, National Institutes for Food and Drug Control. HBSS1 recombinant protein (containing S (1-223 aa) and PreS1 (21-47 aa)) was from Wenjie Tan, Chinese Center for Disease Control and Prevention. NP-KLH (Santa Cruz, sc-396216), NP-BSA (Biosearch Technologies, N-5050H-10), Reactive Oxygen Species Assay Kit (Solarbio, CA1410) was purchased as indicated.

**Virus infection**

The influenza virus A/WSN/1933 (H1N1) strain was from Wenjun Liu, Institute of Microbiology, CAS, China. A/PR8/34 (H1N1) strain was from Yonghui Zhang, Tsinghua University. VACV (Western Reserve strain), H3N2 subtype A/Jiangxi/2005, and H9N2 subtype A/Chicken/Liaoning/1/00 were from Min Fang, Institute of Microbiology, CAS. HSV-1 (Wildtype F strain) and VSV (Indiana strain) were from Hongbing Shu, Wuhan University.

For mouse survival experiments, 6-8 week aged mice were infected with lethal titers of VSV ($5 \times 10^8$ pfu per mouse) intravenously, HSV-1 ($1 \times 10^7$ pfu per mouse) intraperitoneally, VACV ($1 \times 10^7$ pfu per mouse) intranasally, H1N1-PR8 ($1 \times 10^6$ pfu per mouse) intranasally, H1N1-WSN ($1 \times 10^6$ pfu per mouse) intranasally and H3N2 ($1 \times 10^6$ pfu per mouse) intranasally after immunization.

For plaque assay, VSV, HSV-1 and VACV titers were measured by BHK21 cells; influenza virus titers were measured by MDCK cells. BHK21 cells monolayers (100% confluence in 24-well plates) were washed with phosphate-buffered NS (PBS) and infected with different dilutions of virus for 1 h at 37 ℃. Then virus inoculums were removed and washed with PBS twice. Cell monolayers were overlaid with 0.5% methylcellulose in FBS-free DMEM. 48-72 h later, cells were fixed with 0.5% glutaraldehyde and stained with 1% crystal violet dissolved in 70% ethanol. MDCK cell monolayers (100% confluence in 12-well plates) were infected as BHK21 cells, and were overlaid with agar overlay medium (DMEM containing 1% low-melting -point agarose and 1 μg/ml TPCK-treated trypsin) and incubated at 37 ℃ for 48-72 h. Plaques were counted to determine virus titers.

For inactivated virus preparation, different viruses were purified by adding 4% PEG-6000 and 2% NaCl to settle overnight at 4 ℃. Then, viruses were centrifuged at 8 000g for 2 h and precipitates were dissolved in PBS. The purified viruses were inactivated by adding 0.1% formaldehyde and rotated overnight at 37 ℃. After that, viruses were confirmed to be completely inactivated by plaque assay.

**Clinical scoring of IAV model**

The scoring system is based on coat condition, posture and activity according to previously described [2]. Ruffled fur (absent 0, mild 1, severe 2), hunched back (absent 0, mild 1, severe 2) and activity (normal 0, reduced 1, severely reduced 2). The final score = ruffled fur + hunched back + activity.

#### Generation and purification of IAVs

A/PR8/34 (H1N1), A/WSN/1933 (H1N1) A/Jiangxi/262/2005 (H3N2) and H9N2 subtype A/Chicken/Liaoning/1/00 viruses were generated in 10-day-old embryonated chicken eggs for 2 days at 37 ℃. The allantois fluids were collected. Then, the influenza viruses were purified by sucrose density gradient centrifugation with 20 000 g for 2 h at 4 ℃. Finally, viruses were dissolved in PBS and titers were determined by plaque assay.

#### Type I-IFN bioassay

Type I-IFN concentration was measured as previously described [3]. Briefly, an IFN-sensitive luciferase vector was constructed by cloning IFN-stimulated response element (ISRE) into pGL3-Basic Vector (Promega), and was stably transfected into L929 or HT1080 cells. L929-ISRE or HT1080-ISRE [4] cells were seeded to 96-well plates and incubated with mouse or human cell culture supernatants. Recombinant mouse or human IFNβ (R&D Systems) was used as standards. 4 h later, cells were lysed and measured by Luciferase Reporter Assay System (Promega).

#### Inflammasome activation

$1 \times 10^6$ cells were plated in 12-well plate overnight and the medium was changed to opti-MEM with LPS (500 ng/ml) for 3 h. After that, the cells were treated with or without inhibitors for 1 h. Then cells were stimulated with indicated concentrations of $Mn^{2+}$, Alum, ATP (Amresco, Cat# Amresco 0220), VACV (Western Reserve-Vvt7 strain), Salmonella, $Ca^{2+}$ or Silica.

#### Mitochondrial DNA depletion

THP1 cells were cultured in RPMI (GIBCO) supplemented with 10% FBS (GIBCO), 2 mM L-glutamine, 100 μg/ml sodium pyruvate, 50 μg/ml uridine and 50 ng/ml ethidium bromide for 6 days as previously described [5]. Depletion of mtDNA was measured by Real-Time PCR of mtDNA versus genomic DNA.

**Quantitative reverse transcription PCR analysis**

Total RNA was isolated using TRIzol reagent (Invitrogen), according to the manufacturer’s instruction. One microgram of total RNA was converted into cDNA with Oligo dT primer and RevertAid reverse transcriptase (Thermo Scientific). PCR was performed with gene-specific primer sets (Tables S3). Quantitative real-time PCR was performed with Sybr green incorporation on the LightCycler® 96 System (Roche), and the data were presented as mRNA accumulation index ($2^{\Delta\Delta Ct}$).

**Protein expression and purification**

HA1 (11–324 aa) of A/PR8/34(H1N1) was subcloned into pET-28a vector. The recombinant HIS6-HA1 was expressed in *E. coli* BL21 (DE3) with 1 mM IPTG induction for 5 h at 37 ℃. The protein was purified according to the published procedures [6].

**Ultra-structure observation**

MnJ was mounted on 230-mesh copper grids, which were cleaned with dd$H_2O$ for 3 times. Afterwards, the grids were dried overnight and observed under TEM (Ht-7700, Hitachi).

**Mouse immunization**

Mice were immunized with a prime-boost strategy. For intramuscular immunization, each mouse was immunized with 10 μg OVA (InvivoGen, Cat# vac-pova) alone or with MnJ, Imject® Alum (Thermo), Alhydrogel® adjuvant 2% (InvivoGen), Adju-Phos® adjuvant (InvivoGen), Freund's Adjuvant, Complete (Sigma), Freund's Adjuvant, Incomplete (Sigma), SIGMA ADJUVANT SYSTEM (Sigma), Polyethylenimine (Polysciences), MF59 (gift from Dr. Yonghui Zhang) suspended in PBS or normal saline with a final volume 100 μl. Then the mice were boosted on day 7 and 14. For intranasal immunization, each mouse was immunized with 10 μg OVA alone or with MnJ, Cholera Toxin B subunit (Sigma) suspended in PBS or normal saline with a final volume 20 μl after anesthetization. Then the mice were boosted on day 7 and 14.

**Measurement of microelement**

About 200 mg tissue samples were digested with 2 ml $HNO_3$ in a microwave digestion system. Then the amounts of Mn and Al were measured by Inductively Couples Plasma Mass Spectrometry (ICP-MS, Thermo X SERIES II) as previously described [1].

**Antibody titer test**

Serum was collected from whole blood by centrifugation. Bronchoalveolar lavage fluid was collected with 500 μl PBS from lung. Oral lavage fluid was collected with 200 μl PBS from oral cavity. ELISA plated were coated with 2 μg/ml antigens (OVA, HA1, NP-BSA, EV71, HBsAg or HBSS1) in PBS at 4 ºC overnight. After washing with 200 μl Wash Buffer (0.05% Tween20 in PBS) for 5 times, plates were blocked with 200 μl Block Buffer (2 % BSA in PBS) for 2 h at 37 ºC. After washing for 5 times, 100 μl diluted samples were added in plates and incubated for 2 h at 37 ºC.

After washing for 5 times, plates were incubated with HRP-labeled antibodies for 2 h at 37 ℃. After washing for 5 times, plates were incubated with 100 μl 1 × TMB Solution (eBioscience) for 30 min at 37 ºC, followed by adding 100 μl Stop Solution (MultiSciences). Optical density (OD) was read at 450 nm and 570 nm by FlexStation 3 (Molecular Devices). Substrate readings at 570 nm from the readings at 450 nm. Antibody titers were obtained by plotting the maximum serum dilution that gave an optical density > 2 X background.

**_In vitro_ BMDC stimulation**

Immature BMDCs ($1 \times 10^6$) were planted onto 12 well plates, then treated with indicated concentrations of $MnCl_2$ or LPS for 20 h. BMDCs were stained with anti-CD86, anti-CD80 and anti-CD40 (Biolegend). Cell-surface co-stimulatory molecule CD86, CD80 and CD40 were analyzed by FACS.

**_In vivo_ T cell proliferation**

OT-I $CD8^+$ T cells or OT-II $CD4^+$ T cells were isolated from $CD45.1^+$ OT-I or OT-II mice by Mouse CD8 or CD4 T Cell Isolation Kit (Biolegend). CFSE (1 μM)-labelled $CD45.1^+$ OT-I $CD8^+$ or OT-II $CD4^+$ T cells ($2 \times 10^6$) were transferred intravenously into naive $CD45.2^+$ mice before immunization the next day. Inguinal lymph nodes of vaccinated mice were obtained 3 days after immunization and divided $CD4^+$ $CD45.1^+$ or $CD8^+$ $CD45.1^+$ T cells were analyzed by FACS using Precision Count Beads (Biolegend, Cat# 424902).

**_In vivo_ antigen uptake and Mo-DCs accumulation analysis**

C57BL/6 mice were vaccinated in both inguinal regions subcutaneously with PBS, PE, PE+MnJ or PE+Alum. Inguinal dLNs were collected 12 or 24 hours after the immunization. dLN cells were labelled with DAPI, anti-CD11c (Biolegend, Cat#117325), anti-F4/80 (Biolegend, Cat#123115), anti-CD8a (Biolegend, Cat#100721), anti-CD64 (Biolegend, Cat#139315) to identify APCs. Ratio of $PE^+$ APCs ($CD11c^+$ or $F4/80^+$ cells) from dLN cells and ratio of Mo-DCs ($CD64^+$ $CD8a^+$) in APCs ($CD11c^+$ or $F4/80^+$ cells) were analyzed by FACS.

##### *In vivo* cytotoxicity assay

C57BL/6 mice were vaccinated with OVA (10 μg), OVA (10 μg) + MnJ (20 μg) or OVA (10 μg) + Alum (1320 μg) on day 0, 7 and 14. On day 21, vaccinated animals were intravenously injected with $2 \times 10^6$ donor splenocytes from naive C57BL/6 mice. A half was labeled with 0.5 μM CFSE (Biolegend, Cat# 423801) and the other half was labeled with 5 μM CFSE for 10 min at 37 °C. The stained cells at high CFSE concentration were pulsed with 10 μg/ml SIINFEKL peptide for 90 min at 37 ℃ [7]. 36 h after transfer, splenocytes were collected and the specific killing was defined as the percentage of specific lysis = (transferred ratio – experimental ratio) × 100%.

##### Germinal center analysis

Inguinal dLNs were collected 7 days after immunization. LN single cells were stained by FITC labelled anti-CD3 (Biolegend, Cat#100203), APC labelled anti-CD4 (Biolegend, Cat#100411), PE labelled anti-PD1 (Biolegend, Cat#135205), PECy7 labelled anti-CXCR5 (Biolegend, Cat#145515) to analyze Tfh cells or Alexa Fluor® 700 labelled anti-CD45 (Biolegend, Cat#103127), FITC labelled anti-B220 (Biolegend, Cat#103205), APC lebelled anti-GL7 (Biolegend, Cat#144617), PE labelled anti-Fas (Biolegend, Cat#152607) to analyze GC B cells.

**Tumor model**

For subcutaneous tumor, C57BL/6 mice were vaccinated with OVA (10 μg), OVA (10 μg) + MnJ (20 μg) or OVA (10 μg) + Alum (1320 μg) on day 0, 7 and 14. On day 21, vaccinated animals were inoculated subcutaneously on the right hind flank with $3 \times 10^5$ B16-F10-OVA cells. Tumor sizes were measured every 2 days with electronic calipers and calculated by length (mm) × width (mm) × width (mm)/2. Mice with tumors larger than 20 mm on the longest axis were euthanized. IVIS images were captured with a Caliper IVIS Lumina II (Caliper Life Sciences) instrument following light anesthesia with pentobarbital sodium, and intraperitoneal injection of D-luciferin (Cayman Chemical, 14681) (0.5 mg/g per mouse). Images were quantified using Living Image 4.0.

For metastatic tumor, C57BL/6 mice were vaccinated with OVA (10 μg) or OVA (10 μg) + MnJ (20 μg) on day 0, 7 and 14. On day 21, vaccinated animals were inoculated intravenously with $3 \times 10^5$ B16-F10-OVA-GFP cells. Mice were euthanized 20 days after inoculation. The images of lungs and HE-stained lung sections were recorded.

**Tumor infiltrating T cell analysis**

Tumor tissues were collected on day 14 after inoculation. For FACS analysis, the tissues were cut into pieces and incubated in a PBS solution containing Collagenase A (0.3 mg/ml) and DNase I (0.01 mg/ml) for 60 min at 37 ℃ under gentle rotation. Digestion was stopped by adding FBS on ice. The supernatants were centrifuged at 4000 rpm for 10 min at 4 ℃. The samples were then resuspended in 1 ml PBS and filtered through a 100 mesh nylon sieve, followed by washing. Red blood cells were removed by ACK lysis buffer (155 mM $NH_4Cl$, 10 mM $KHCO_3$, 0.1 mM EDTA). Cells were incubated with APC labeled anti-CD4 (Biolegend, Cat# 100411) and PE/Cy7 labeled anti-CD8a (Biolegend, Cat# 100721). Tumor infiltrating T cells were analyzed by FACS. Fixed tumor tissues were embedded in 4% paraffin and cut into 4 μm sections. $CD4^+$ T cells were stained with anti-CD4 monoclonal antibody (Servicebio, GB13064-2) and FITC-labeled Goat Anti-Rabbit IgG (H+L) (Servicebio, GB22303). $CD8^+$ T cells were stained with anti-CD8 monoclonal antibody (Servicebio, GB11068) and Cy3 conjugated Goat Anti-rabbit IgG (H+L) (Servicebio, GB21303). Tissues were stained with DAPI (Servicebio, G1012) finally. Images were acquired by using a confocal microscope (Andor Dragonfly).

**Tetramer staining**

C57BL/6 mice were immunized with OVA (100 μg) and OVA (100 μg) + $Mn^{2+}$ (20 μg) on day 0, 7 and 14. On day 21, splenocytes ($1 \times 10^6$) from immunized mice were isolated and hemolyzed with ACK lysis buffer. Cells were incubated with PE labeled H2-Kb/OVA (SIINFEKL) tetramers (MBL, TS-5001-1C) for 30 min at 4 ℃. Then add PE/Cy7 labeled anti-CD8a (Biolegend, Cat#100721) to incubate for 30 min at 4 ℃. After washing, cells were analyzed by FACS.

**Statistical analysis**

Student’s t-test was used to analyze data. Survival curves were compared using Mantel-Cox test.

All results are expressed as the mean ± SEM. Unpaired two-tailed t tests were used for comparison of two groups. A two-way ANOVA was performed when both time and treatment were compared. For the survival studies, a Log-Rank (Mantel-Cox) test was used. $p < 0.05$ was considered statistically-significant and denoted as follows: $*p < 0.05$, $**p < 0.01$, $***p < 0.001$, $****p < 0.0001$. Statistical analyses were performed by using GraphPad PRISM.

**Data and software availability**

RNA-seq data have been deposited in Gene Expression Omnibus under accession no. GSE126586.

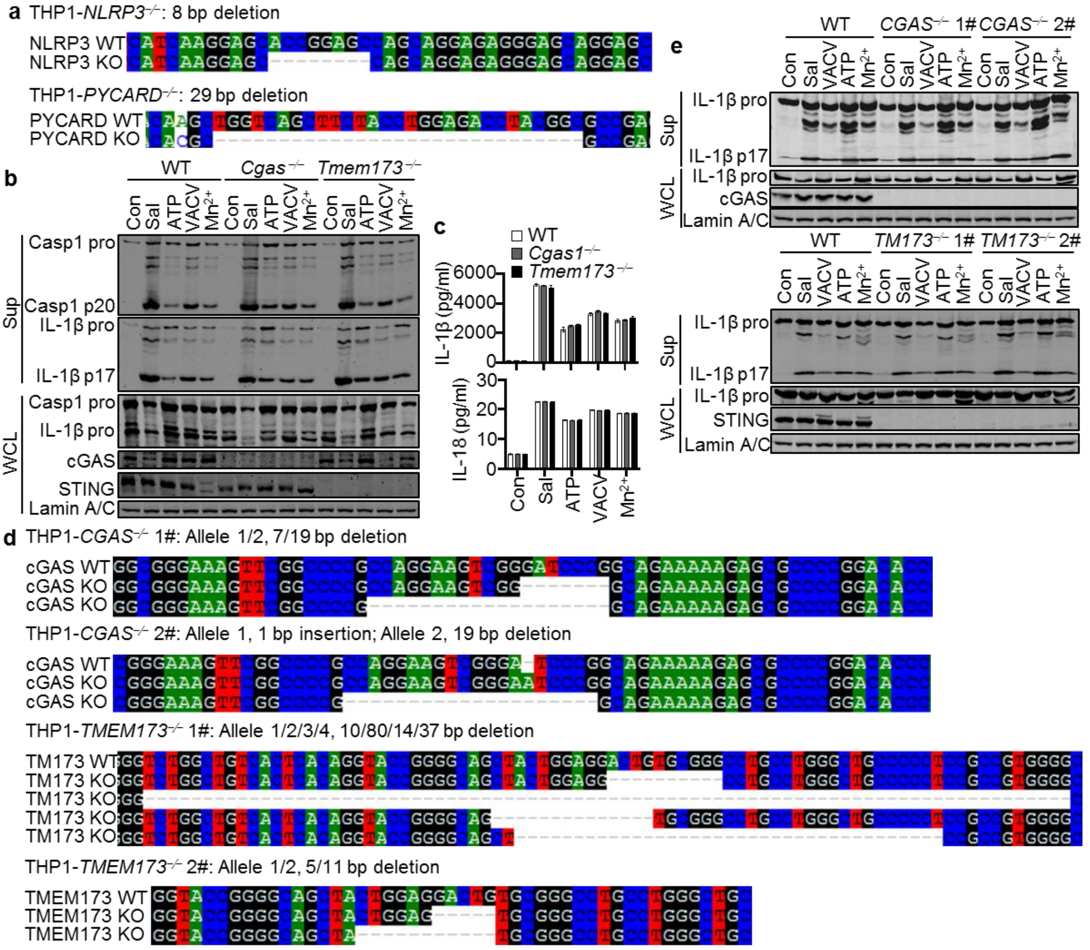


**Extended Data Fig. 1| $Mn^{2+}$ Activates NLRP3 independent of cGAS or STING. a,** Genotyping of *NLRP3*−/− and *PYCARD*−/− THP1 cells. **b, c,** LPS-primed WT, *Cgas*−/− and *Tmem173*−/− peritoneal macrophages were stimulated with Salmonella (MOI = 10) or ATP (5 mM) for 1 h, VACV (MOI = 10) or $Mn^{2+}$ (200 μM) for 5 h. Supernatants (Sup) and whole cell lysates (WCL) were analyzed by immunoblotting with the indicated antibodies (**b**). IL-1β and IL-18 in supernatants were analyzed by ELISA (**c**). Sal, Salmonella. **d,** Genotyping of *cGAS*−/− and *TMEM173*−/− THP1 cells. **e,** LPS-primed WT, *cGAS*−/− or *TMEM173*−/− THP1 cells were treated with Salmonella (MOI = 10), ATP (5 mM) and $MnCl_2$ (1 mM). The cleavage of IL-1β was analyzed by Western blot. One representative experiment of at least three independent experiments is shown, and each was done in triplicate.

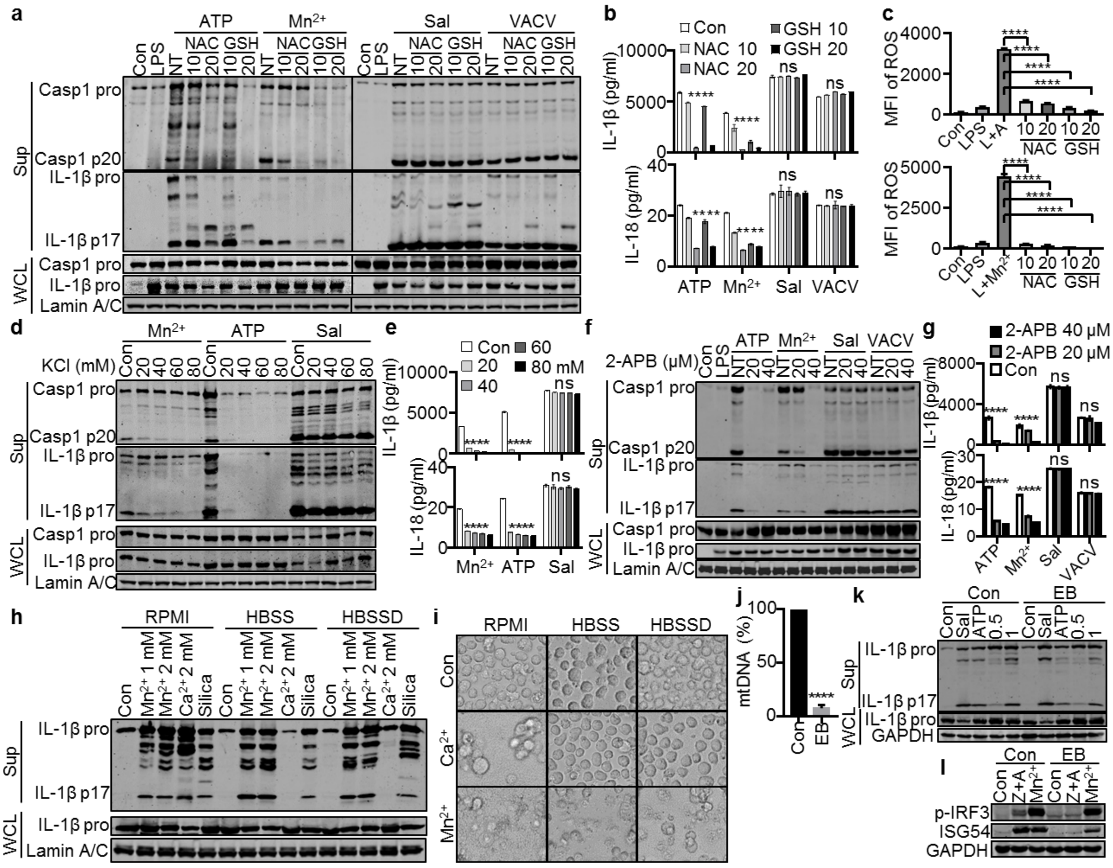


**Extended Data Fig. 2| ROS, $K^+$ efflux and $Ca^{2+}$ Release Are Required for $Mn^{2+}$-activated Inflammasome. a, b,** Western blot (**a**) and ELISA analysis (**b**) of inflammasome activation of LPS-primed peritoneal macrophages treated with ATP (5 mM), $MnCl_2$ (0.5 mM), Salmonella (MOI = 10) or VACV (MOI = 10), which were pretreated with ROS inhibitor NAC (10 or 20 mM) or GSH (10 or 20 mM) for 1 h. **c,** Intracellular ROS levels were determined by FACS using DCFH-DA ROS probe. L + A, LPS + ATP. L + $Mn^{2+}$, LPS + $Mn^{2+}$. **d, e,** Western blot (**d**) and ELISA analysis (**e**) of inflammasome activation of LPS-primed peritoneal macrophages treated with ATP (5 mM), $MnCl_2$ (0.5 mM) or Salmonella (MOI = 10), which were cultured in Opti-MEM with the indicated extra concentrations of extracellular $K^+$. **f, g,** Western blot (**f**) and ELISA analysis (**g**) of inflammasome activation of LPS-primed peritoneal macrophages treated with ATP

(5 mM), $Mn^{2+}$ (0.5 mM), Salmonella (MOI = 10) or VACV (MOI = 10), which were pretreated with $Ca^{2+}$ release inhibitor 2-APB for 1 h. **h,** Western blot analysis of inflammasome activation of LPS-primed THP1 cells treated with $MnCl_2$ (1 and 2 mM), $CaCl_2$ (2 mM) and Silica (0.1 mg/ml) for 5 h cultured in RPMI, HBSS (5.33 mM KCl, 0.44 mM $KH_2PO_4$, 4.17 mM $NaHCO_3$, 137.93 mM NaCl, 0.34 mM $Na_2HPO_4$, 5.56 mM D-Glucose) or HBSSD (5.33 mM KCl, 137.93 mM NaCl, 5.56 mM D-Glucose). **i,** Images of pyroptosis of THP1 cells treated in (**h**). **j,** Quantitative PCR analysis of mtDNA versus genomic DNA from control and EB-co-cultured THP1 cells (n = 3). **k,** Western blot analysis of inflammasome activation of LPS-primed control and EB-co-cultured THP1 cells treated with Salmonella (MOI = 10), ATP (5 mM) and $MnCl_2$ (0.5 and 1 mM). **l,** Western blot analysis of Type I-IFN production in control and EB-co-cultured THP1 cells treated with Z-VAD-FMK (10 μM) + ABT263 (10 μM) or $MnCl_2$ (0.5 mM) for 18 h. Z+A, Z-VAD-FMK + ABT263. One representative experiment of at least three independent experiments is shown, and each was done in triplicate. Error bars represent SEM; **b, c, e, g, j,** data were analyzed by an unpaired t test. ns, not significant; * $P < 0.05$; ** $P < 0.01$; *** $P<0.001$; **** $P < 0.0001$.

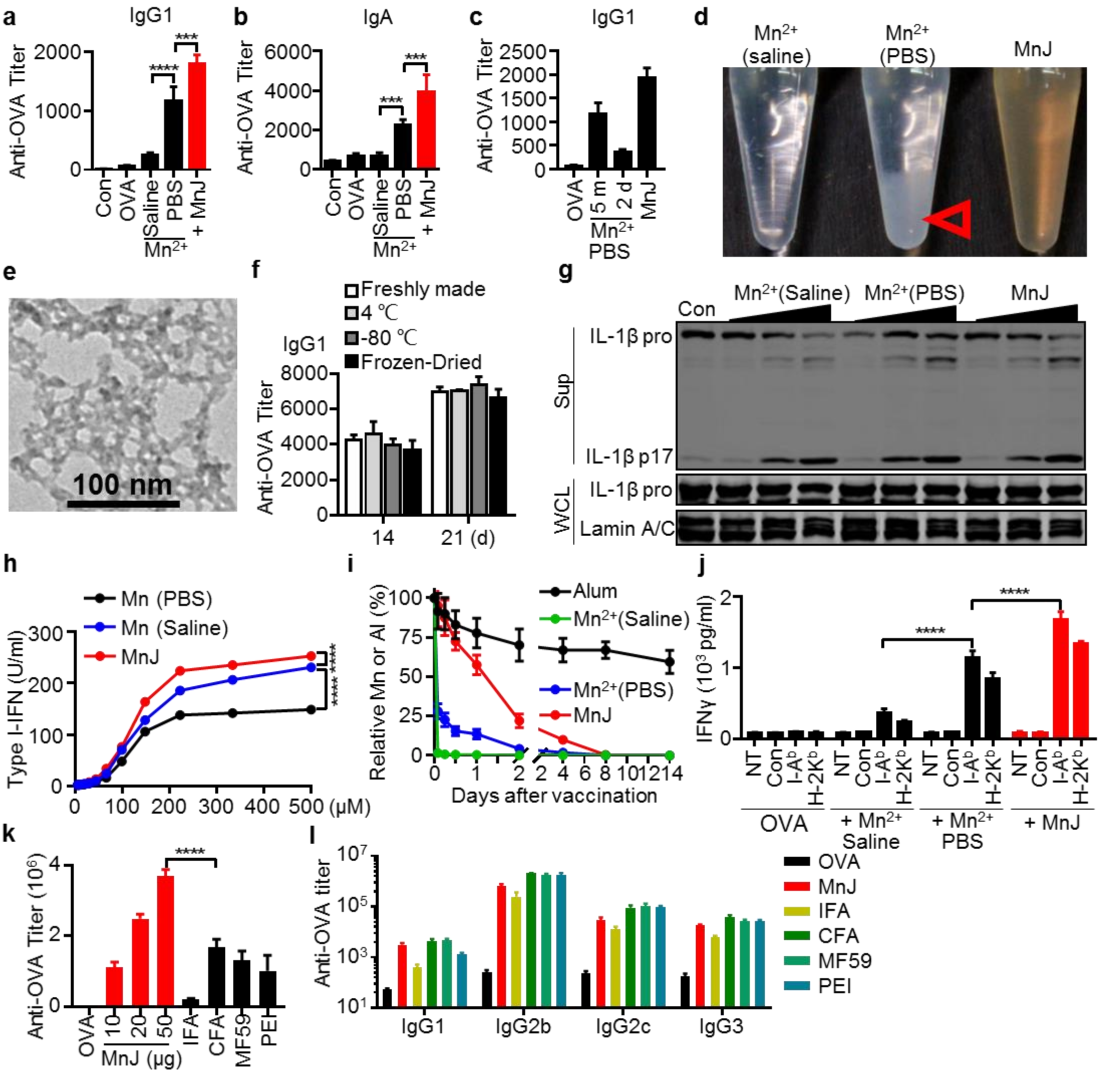


**Extended Data Fig. 3| Properties of $Mn_2OHPO_4$ (MnJ) Adjuvant. a,** The WT mice were immunized intramuscularly with PBS, OVA (10 μg), OVA (10 μg) + $Mn^{2+}$ (10 μg in normal saline), OVA (10 μg) + $Mn^{2+}$ (10 μg in PBS) or OVA (10 μg) + MnJ (10 μg in normal saline) on day 0, 7 and 14. Sera were collected on day 21 to measure OVA-specific IgG1 by ELISA (n = 3). **b,** The WT mice were immunized intranasally with PBS, OVA (10 μg), OVA (10 μg) + $Mn^{2+}$ (5 μg in normal saline), OVA (10 μg) + $Mn^{2+}$ (5 μg in PBS) or OVA (10 μg) + MnJ (5 μg in normal saline) on day 0, 7 and 14. Sera were collected on day 21 to measure OVA-specific IgA by ELISA (n = 3). **c,** The WT mice were immunized intramuscularly with OVA (10 μg), OVA (10 μg) + $Mn^{2+}$ (10 μg in PBS for 5 min), OVA (10 μg) + $Mn^{2+}$ (10 μg in PBS for 2 d) or OVA (10 μg) + MnJ (10 μg in normal saline) on day 0, 7 and 14. Sera were collected on day 21 to measure OVA-specific IgG1 by ELISA (n = 3). **d,** $MnCl_2$ (20 mM each) in normal saline, PBS or saline containing 25 mM $Na_3PO_4$ (MnJ), mixtures were settled overnight before pictures were taken. Aggregated and precipitated Mn salts in PBS was indicated by an open arrow. **e,** Representative image of MnJ nanoparticles. Images were obtained by TEM. Scale bar = 100 nm. **f,** The WT mice were immunized intramuscularly with OVA (10 μg) + MnJ (10 μg freshly made), OVA (10 μg) + MnJ (10 μg kept at 4 ℃), OVA (10 μg) + MnJ (10 μg kept at -80 ℃) or OVA (10 μg) + MnJ (10 μg frozen-dried) on day 0, 7 and 14. Sera were collected on day 21 to measure OVA-specific IgG1 by ELISA (n = 8). **g,** LPS-primed THP1 cells in HBSSD (5.33 mM KCl, 137.93 mM NaCl, 5.56 mM D-Glucose) were treated with $MnCl_2$ + normal saline, $MnCl_2$ + PBS or MnJ for 5 h. The cleavage of IL-1β was measured by Western blot. All $Mn^{2+}$ stock solutions contain 20 mM $Mn^{2+}$ and cells were treated with 250, 500 and 1000 μM $Mn^{2+}$. **h,** THP1 cells were treated with $MnCl_2$ + normal saline, $MnCl_2$ + PBS or MnJ for 24 h. All $Mn^{2+}$ stock solutions contain 20 mM $Mn^{2+}$ and cells were treated with 6, 9, 13, 20, 29, 44, 66, 99, 148, 222, 333 and 500 μM $Mn^{2+}$. The Production of Type I-IFN was measured by bioassay. **i,** The WT mice were injected intramuscularly with 100 μg Alum, 20 μg $MnCl_2$ (in normal saline), 20 μg $MnCl_2$ (in PBS) or 20 μg MnJ. Muscles at the injection site (100 mg) were collected at the indicated times to measure the amounts of Mn and Al by ICP-MS (n = 3). **j,** The WT mice were injected intramuscularly with OVA (10 μg), OVA (10 μg) + $Mn^{2+}$ (10 μg in normal saline), OVA (10 μg) + $Mn^{2+}$ (10 μg in PBS) or OVA (10 μg) + MnJ (10 μg in normal saline) on day 0, 7 and 14. Splenocytes were collected on day 21, and stimulated with

OVA peptides. IFNγ secreted by T cells was measured by ELISA. **k, l,** OVA-specific total IgG, IgG1, IgG2b, IgG2c, IgG3 were measured by ELISA on day 21 after immunization with OVA (10 μg), OVA (10 μg) + indicated amounts of MnJ, IFA (50 μl), CFA (50 μl), MF59 (50 μl) or PEI (100 μg) intramuscularly on day 0, 7 and 14 ($n = 3$). One representative experiment of at least three independent experiments is shown, and each was done in triplicate. Error bars represent SEM; **a, b, j, k,** data were analyzed by an unpaired t test; **h,** data were analyzed by two-way ANOVA. ns, not significant; * $P < 0.05$; ** $P < 0.01$; *** $P<0.001$; **** $P < 0.0001$.

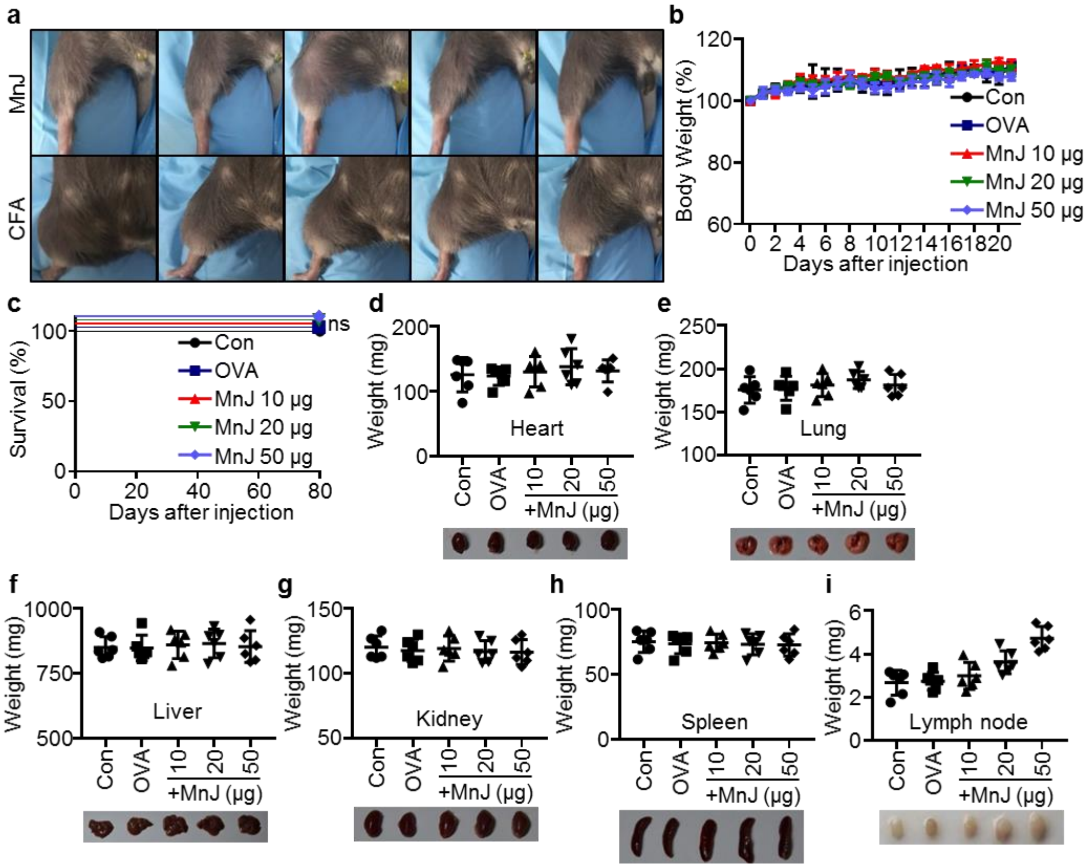


**Extended Data Fig. 4| MnJ Does Not Show Visible Toxicity. a,** Swelling and granulomas formed at the site of injection from mice (n = 5) immunized intramuscularly with CFA (50 µl CFA + 50 µl normal saline) after one injection (bottom) or mice were immunized intramuscularly with 50 µg MnJ (in 100 µl normal saline, once a week) after three injections (Top). Photos were taken on day 7 (CFA) or day 21 (MnJ) respectively. **b, c,** The WT mice were immunized intramuscularly with OVA (10 µg) + MnJ (10, 20 and 50 µg) on day 0, 7 and 14. Their body weight (**b**) and survival (**c**) were monitored for 3 weeks. **d-i,** On day 40, hearts (**d**), lungs (**e**), livers (**f**), kidneys (**g**), spleens (**h**) and inguinal lymph nodes (**i**) were collected and organ weights were recorded (n = 6). One representative experiment of at least three independent experiments is shown, and each was done in triplicate. Error bars represent SEM; **c,** survival plot data were analyzed with log-rank (Mantel–Cox) tests. ns, not significant.

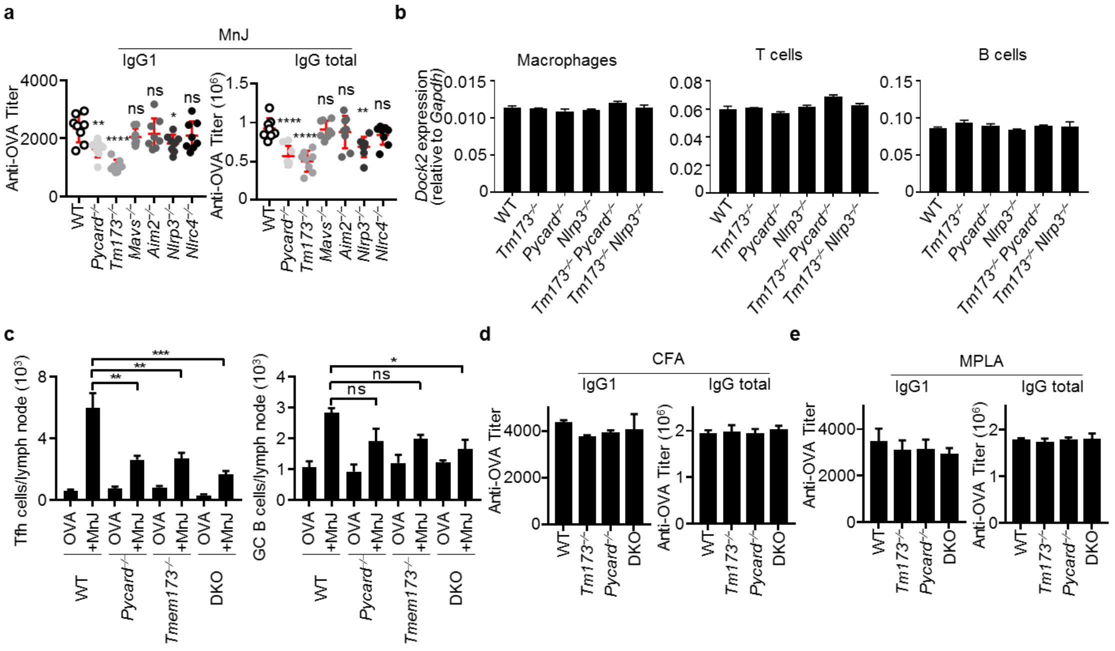


**Extended Data Fig. 5| Both cGAS-STING and NLRP3 Inflammasome Contribute to Adjuvant Activity of MnJ. a,** The WT, *Pycard*$^{-/-}$, *Tmem173*$^{-/-}$, *Mavs*$^{-/-}$, *Aim2*$^{-/-}$, *Nlrp3*$^{-/-}$ and *Nlrc4*$^{-/-}$ mice were immunized intramuscularly with OVA (10 μg) + MnJ (10 μg) on day 0, 7 and 14. Sera were collected on day 21 to measure OVA-specific IgG1 and IgG total by ELISA (n = 8). **b,** Quantitative RT-PCR analysis of *Dock2* expression in the peritoneal macrophages, T cells and B cells of WT, *Tmem173*$^{-/-}$, *Pycard*$^{-/-}$, *Nlrp3*$^{-/-}$, *Tmem173*$^{-/-}$ *Pycard*$^{-/-}$ and *Tmem173*$^{-/-}$ *Nlrp3*$^{-/-}$ mice. **c,** Numbers of Tfh or GC B cells in dLN from WT, *Pycard*$^{-/-}$, *Tmem173*$^{-/-}$, DKO mice were analyzed by FACS. Live cells were identified by DAPI staining. Among live singlet cells, CD4$^{+}$ T cells were identified as the cell subset double positive for CD3 and CD4. Among CD4$^{+}$ T cells, Tfh cells were identified as PD1$^{+}$ CXCR5$^{+}$ cells. B cells were identified as the cell subset double positive for CD45 and B220. Among B cells, GC B cells were identified as Fas$^{+}$ GL7$^{+}$ cells. **d, e,** The WT, *Tmem173*$^{-/-}$, *Pycard*$^{-/-}$ and DKO mice were immunized intramuscularly with OVA (10 μg) + CFA (50 μl) (**c**) or MPLA (2 μl) (**d**) on day 0, 7 and 14. Sera were collected on day 21 to measure OVA-specific IgG1 and IgG total by ELISA (n = 3). MPLA was SIGMA ADJUVANT SYSTEM (R) (S6322-1VL). One representative experiment of at least three independent experiments is shown, and each was done in triplicate. Error bars represent SEM; **a,c,** data were analyzed by an unpaired t test. ns, not significant; * $P < 0.05$; ** $P < 0.01$; *** $P<0.001$; **** $P < 0.0001$.

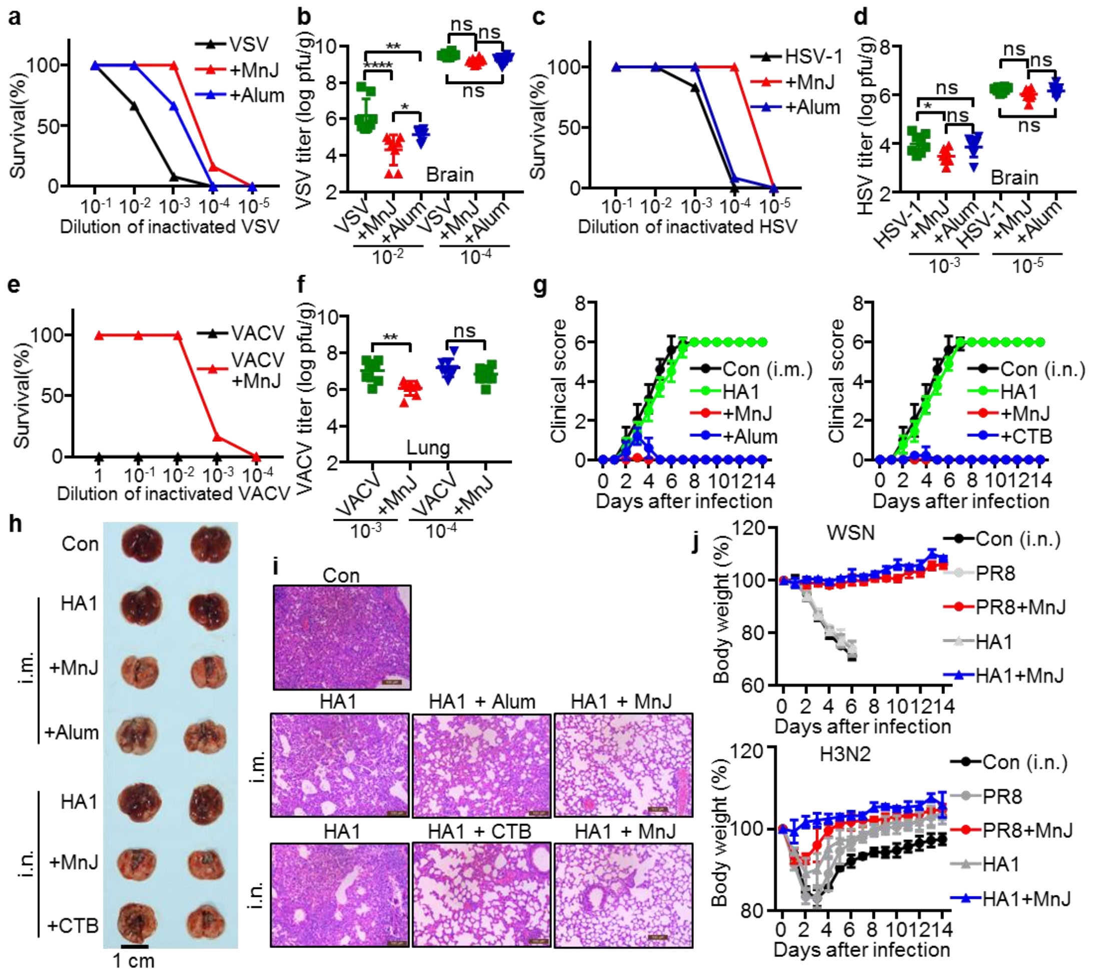


**Extended Data Fig. 6| MnJ Shows Strong Adjuvant Effects against Viruses. a, b,** The WT mice were immunized intramuscularly with PBS, inactivated $10^{-1}$ VSV ($1 \times 10^7$ pfu), $10^{-2}$ VSV ($1 \times 10^6$ pfu), $10^{-3}$ VSV ($1 \times 10^5$ pfu), $10^{-4}$ VSV ($1 \times 10^4$ pfu) or $10^{-5}$ VSV ($1 \times 10^3$ pfu) with or without MnJ (10 μg) and Alum (1320 μg) on day 0. On day 10, these mice were infected intravenously with a lethal dose of VSV. The survival was monitored for 2 weeks (n = 12) (**a**). Viral loads in brain were measured 5 days after infection (n = 8) (**b**). **c, d,** The WT mice were immunized intramuscularly with PBS, inactivated $10^{-1}$ HSV-1 ($1 \times 10^6$ pfu), $10^{-2}$ HSV-1 ($1 \times 10^5$ pfu), $10^{-3}$ HSV-1 ($1 \times 10^4$ pfu), $10^{-4}$ HSV-1 ($1 \times 10^3$ pfu) or $10^{-5}$ HSV-1 ($1 \times 10^2$ pfu) with or without MnJ (10 μg) and Alum (1320 μg) on day 0. On day 10, these mice were infected intraperitoneally with a lethal dose of HSV-1. The survival was monitored for 2 weeks (n = 12) (**c**). Viral loads in brain were measured 5 days after infection (n = 8) (**d**). **e, f,** The WT mice were immunized intranasally with PBS, inactivated VACV ($2 \times 10^6$ pfu), $10^{-1}$ VACV ($2 \times 10^5$ pfu), $10^{-2}$ VACV ($2 \times 10^4$ pfu), $10^{-3}$ VACV ($2 \times 10^3$ pfu) or $10^{-4}$ VACV ($2 \times 10^2$ pfu) with or without MnJ (5 μg) on day 0 and 7. On day 14, these mice were infected intranasally with a lethal dose of VACV. The survival was monitored for 2 weeks (n = 12) (**e**). Viral loads in lung were measured 5 days after infection (n = 8) (**f**). **g,** Clinical scores of mice in Fig. 4G were daily evaluated for 2 weeks.

**h,** Images of lungs from mice treated in Fig. 4G were recorded on day 5. **i,** HE-stained lung sections from mice in Fig. 4G were analyzed on day 5 by Leica CTR5000. **j,** The WT mice were immunized intranasally with PBS, inactivated PR8 ($5 \times 10^6$ pfu), inactivated PR8 ($5 \times 10^6$ pfu) + MnJ (10 μg) for 2 times or HA1 (5 μg) or HA1 (5 μg) + MnJ (10 μg) for 3 times. Then, these mice were infected intranasally with a lethal dose of H1N1 WSN subtype A/WSN/1933 and H3N2 subtype A/Jiangxi/262/2005. Body weight was recorded for 2 weeks (n = 5). One representative experiment of at least three independent experiments is shown, and each was done in triplicate. Error bars represent SEM; **b, d, f,** data were analyzed by an unpaired t test. ns, not significant; * $P < 0.05$; ** $P < 0.01$; *** $P<0.001$; **** $P < 0.0001$.

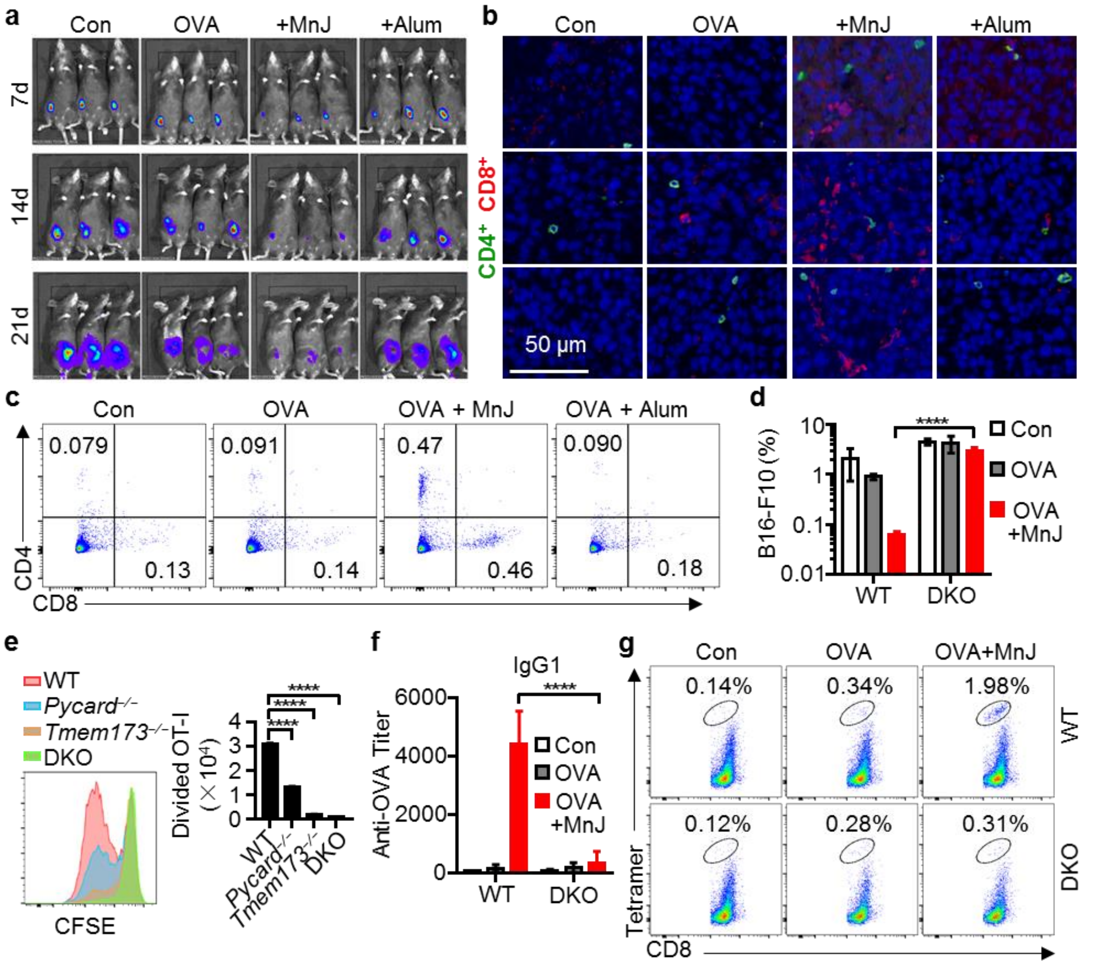


**Extended Data Fig. 7| MnJ Greatly Potentiates Antitumor Effect. a-c,** The WT mice were immunized intramuscularly with PBS, OVA (10 μg), OVA (10 μg) + MnJ (20 μg) or OVA (10 μg) + Alum (1320 μg) on day 0, 7 and 14. On day 21, these mice were inoculated with B16-OVA-Fluc cells (3 × $10^5$) subcutaneously. Representative IVIS images were acquired on day 7, 14 and 21 after inoculation (**a**). Some tumors were collected on day 14 and immune-staining of $CD4^+$ and $CD8^+$ T cells in tumor sections were recorded (**b**). Tumor infiltrating T cells were analyzed by FACS on day 14 (**c**). **d,** The WT and DKO mice were immunized intramuscularly with PBS, OVA (10 μg), or OVA (10 μg) + MnJ (20 μg) on day 0, 7 and 14. On day 21, these mice were inoculated with B16-F10-OVA-GFP (3 × $10^5$) intravenously. Percentage of $GFP^+$ B16-F10 in lung was analyzed by FACS on day 20 after inoculation. **e,** CD45.1$^+$ OT-I CD8$^+$ T cells were labeled with CFSE and transferred to CD45.2$^+$ WT, *Tmem173*$^{-/-}$, *Pycard*$^{-/-}$ and DKO mice. These mice were then immunized with OVA (1 μg) + MnJ (10 μg). After 3 days, T cell proliferation was analyzed by FACS (n = 3). **f,** OVA-specific IgG1 in serum from mice in (**d**) was measured by ELISA on day 20. **g,** The WT and *Tmem173*$^{-/-}$*Pycard*$^{-/-}$ DKO mice were immunized intramuscularly with PBS, OVA (10 μg), OVA (10 μg) + MnJ (20 μg) on day 0, 7 and 14. On day 21, the percentage of tetramer$^+$ CD8$^+$ T cells in spleens of these mice was analyzed by FACS. One representative experiment of at least three independent experiments is shown, and each was done in triplicate. Error bars represent SEM; **d, e, f,** data were analyzed by using an unpaired t test. **** P < 0.0001.

**Extended Data Table 1| Information of Peripheral Blood Mononuclear Cell Donors.**

| **Number** | **Age** | **Sex** | **Physical condition** |
|---|---|---|---|
| 1 | 26 | Male | Healthy |
| 2 | 22 | Male | Healthy |
| 3 | 22 | Female | Healthy |
| 4 | 25 | Female | Healthy |
| 5 | 23 | Female | Healthy |
| 6 | 21 | Female | Healthy |
| 7 | 24 | Female | Healthy |

**Extended Data Table 2| Target sequences of sgRNAs.**

| Target of sgRNA | Sequence |
|---|---|
| cGAS | 5’-CCGCCAGGAAGTCGGGATCC-3’ |
| STING | 5’-CAGCTACTGGAGGACTGTGC-3’ |
| NLRP3 | 5’-TGCGTCTCATCAAGGAGCAC-3’ |
| PYCARD | 5’-CAAGCTGGTCAGCTTCTACC-3’ |

**Extended Data Table 3| Primers for qRT-PCR.**

| Primers | Forward (5’- 3’) | Reverse (5’- 3’) |
|---|---|---|
| Ifnb1 | GCCTTTGCCATCCAAGAGATGC | ACACTGTCTGCTGGTGGAGTTC |
| Ifna1 | GGATGTGACCTTCCTCAGACTC | ACCTTCTCCTGCGGGAATCCAA |
| Ifna2 | ATCCAGAAGGCTCAAGCCATCC | GGAGGGTTGTATTCCAAGCAGC |
| Ifna4 | GCAATGACCTCCATCAGCAGCT | GTGGAAGTATGTCCTCACAGCC |
| Ifit1 | TACAGGCTGGAGTGTGCTGAGA | CTCCACTTTCAGAGCCTTCGCA |
| Ifit2 | CGAACTACCGTCTGGATGACTG | CTTCAACCAGCGCCATTGCTTG |
| Ifit3 | GCTCAGGCTTACGTTGACAAGG | CTTTAGGCGTGTCCATCCTTCC |
| Ccl4 | ACCCTCCCACTTCCTGCTGTTT | CTGTCTGCCTCTTTTGGTCAGG |
| Ccl5 | CCTGCTGCTTTGCCTACCTCTC | ACACACTTGGCGGTTCCTTCGA |
| Il6 | TACCACTTCACAAGTCGGAGGC | CTGCAAGTGCATCATCGTTGTTC |
| Il10 | CGGGAAGACAATAACTGCACCC | CGGTTAGCAGTATGTTGTCCAGC |
| Il12a | ACGAGAGTTGCCTGGCTACTAG | CCTCATAGATGCTACCAAGGCAC |
| Il1b | TGGACCTTCCAGGATGAGGACA | GTTCATCTCGGAGCCTGTAGTG |
| Il18 | GACAGCCTGTGTTCGAGGATATG | TGTTCTTACAGGAGAGGGTAGAC |
| Isg15 | CATCCTGGTGAGGAACGAAAGG | CTCAGCCAGAACTGGTCTTCGT |
| Rsad2 | GGAAGGTTTTCCAGTGCCTCCT | ACAGGACACCTCTTTGTGACGC |

| | | |
|---|---|---|
| Tnfa | GGTGCCTATGTCTCAGCCTCTT | GCCATAGAACTGATGAGAGGGAG |
| Dock2 | TTGCTCAGCCAGCTACTGTATG | TTGGTGATGACAGGAAGCAGAAT |
| Gapdh | CATCACTGCCACCCAGAAGACTG | ATGCCAGTGAGCTTCCCGTTCAG |
| IL1B | CCACAGACCTTCCAGGAGAATG | GTGCAGTTCAGTGATCGTACAGG |
| IL18 | GATAGCCAGCCTAGAGGTATGG | CCTTGATGTTATCAGGAGGATTCA |
| TNFA | CTCTTCTGCCTGCTGCACTTTG | ATGGGCTACAGGCTTGTCACTC |
| GAPDH | GTCTCCTCTGACTTCAACAGCG | ACCACCCTGTTGCTGTAGCCAA |
| gDNA (GAPDH) | CTGTTCGACAGTCAGCCGCATC | GCGCCCAATACGAC CAAATCCG |
| mtDNA (COXII) | CCCCACATTAGGCTTAAAAACAGAT | TATACCCCCGGTCGTGTAGC |
